# Titan cells formation in *Cryptococcus neoformans* is finely tuned by environmental conditions and modulated by positive and negative genetic regulators

**DOI:** 10.1101/191668

**Authors:** Benjamin Hommel, Liliane Mukaremera, Radames J. B. Cordero, Carolina Coelho, Christopher A. Desjardins, Aude Sturny-Leclère, Guilhem Janbon, John R. Perfect, James A. Fraser, Arturo Casadevall, Christina A. Cuomo, Françoise Dromer, Kirsten Nielsen, Alexandre Alanio

## Abstract

The pathogenic fungus *Cryptococcus neoformans* exhibits morphological changes in cell size during lung infection, producing both typical size 5 to 7 µm cells and large titan cells (> 10 µm and up to 100 µm). We found and optimized *in vitro* conditions that produce titan cells in order to identify the ancestry of titan cells, the environmental determinants, and the key gene regulators of titan cell formation. Titan cells generated *in vitro* harbor the main characteristics of titan cells produced *in vivo* including their large cell size (>10 µm), polyploidy with a single nucleus, large vacuole, dense capsule, and thick cell wall. Here we show titan cells derived from the enlargement of progenitor cells in the population independent of yeast growth rate. Change in the incubation medium, hypoxia, nutrient starvation and low pH were the main factors that trigger titan cell formation, while quorum sensing factors like the initial inoculum concentration, pantothenic acid, and the quorum sensing peptide Qsp1p also impacted titan cell formation. Inhibition of ergosterol, protein and nucleic acid biosynthesis altered titan cell formation, as did serum, phospholipids and anti-capsular antibodies in our settings. We explored genetic factors important for titan cell formation using three approaches. Using H99-derivative strains with natural genetic differences, we showed that titan cell formation was dependent on *LMP1* and *SGF29* genes. By screening a gene deletion collection, we also confirmed that *GPR4/5-RIM101*, and *CAC1* genes were required to generate titan cells and that the *PKR1*, *TSP2*, *USV101* genes negatively regulated titan cell formation. Furthermore, analysis of spontaneous Pkr1 loss-of-function clinical isolates confirmed the important role of the Pkr1 protein as a negative regulator of titan cell formation. Through development of a standardized and robust *in vitro* assay, our results provide new insights into titan cell biogenesis with the identification of multiple important factors/pathways.

**Author Summary:** *Cryptococcus neoformans* is a yeast that is capable of morphological change upon interaction with the host. Particularly, in the lungs of infected mice, a subpopulation of yeast enlarges, producing cells up to 100 µm in cell body diameter – referred to as titan cells. Along with their large size, the titan cells have other unique characteristics such as thickened cell wall, dense capsule, polyploidization, large vacuole with peripheral nucleus and cellular organelles. The generation of a large number of such cells outside the lungs of mice has been described but was not reproducible nor standardized. Here we report standardized, reproducible, robust conditions for generation of titan cells and explored the environmental and genetic factors underlying the genesis of these cells. We showed that titan cells were generated upon stresses such as change in the incubation medium, nutrient deprivation, hypoxia and low pH. Using collections of well characterized reference strains and clinical isolates, we validated with our model that the cAMP/PKA/Rim101 pathway is a major genetic determinant of titan cell formation. This study opens the way for a more comprehensive picture of the ontology of morphological changes in *Cryptococcus neoformans* and its impact on pathobiology of this deadly pathogen.

## Introduction

The ubiquitous environmental yeast *Cryptococcus neoformans* is a basidiomycetous yeast that has been estimated to cause over 200,000 new cases of meningoencephalitis with greater than 180,000 deaths per year worldwide [1], occurring mostly in immunocompromised individuals with acquired immunodeficiency syndrome (AIDS) [2]. The natural history of most of the cases of this invasive fungal infection proceeds through 3 stages: (i) primary infection via inhalation of desiccated yeasts or basidiospores, with development of sub-clinical pneumonia and spontaneous resolution via granuloma formation; (ii) latency of dormant yeast cells, as demonstrated epidemiologically [3] and biologically [4] and (iii) reactivation and dissemination upon immunosuppression, with meningoencephalitis as the most severe clinical presentation of disease [5]. From the environment to interactions with hosts, the yeasts experience drastic changes that reflect a capacity to rapidly adapt and survive in host tissues and cause disease [4,6–9]. In hosts, *C. neoformans* is exposed to various stresses including high temperature, nutrient deprivation, low pH, hypoxia and high levels of free radicals [10].

In response to the host environment, morphological changes are required to survive and cause disease [11]. Specifically, *C. neoformans* alters its morphology and produces enlarged cells referred to as giant or “titan cells” [12, 13]. This phenomenon has been observed in animal and insect models of cryptococcosis, as well as in human lung and brain infections [12–16]. Titan cells have increased cell body size, ranging from 10 µm up to 100 µm in diameter [13,17–21] as compared to the 5-7 µm size of typical cells. Studies exploring titan cell biology have revealed that these are: (i) uninucleate polyploid cells [12,13,20]; (ii) possess a large single vacuole; (iii) are surrounded by a thick cell wall [22]; and (iv) have a dense and highly crosslinked capsule [13,17,22]. Titan cells also exhibit increased resistance to various stresses including phagocytosis [21], oxidative and nitrosative stress [20, 21], and resistance to the antifungal drug fluconazole [20]. Importantly, titan cell production also enhances dissemination, survival and virulence in a mouse model of infection [19]. Titan cell formation is known to be regulated by the G-protein coupled receptors Gpr5 and Ste3a, that signal through the Gα subunit protein Gpa1 to trigger the cyclic adenosine monophosphate / protein kinase A (cAMP/PKA) signaling pathway [15,20,23–26]. The cAMP/PKA pathway is critical for regulation of other virulence factors in *C. neoformans*, including capsule formation [23, 24], notably through its action on the ubiquitin-proteasome pathway [27]. Pka1 is known to be negatively regulated by the protein Pkr1, and *pkr1Δ* mutant strains exhibit enlarged capsule [23]. Further studies show that titan cells formation is increased by high *PKA1* expression or low Pkr1 activity and is decreased by low *PKA1* expression [6]. Downstream of the PKA pathway, Rim101, a major transcription factor that again controls production of many virulence factors, is also necessary for titan cell production [18].

To date, studies of titan cell formation have been hindered by an inability to consistently and reproducibly generate large quantities of titan cells *in vitro*. Although several methods have been reported for inducing large cells *in vitro*, there have been persistent problems in easily and consistently implementing these protocols across laboratories [17], presumably because the variables that contribute to titan cell inducing conditions are not well understood.

In this study, we identified robust *in vitro* conditions that generate enlarged cells with many of the *in vivo* titan cell characteristics and used this protocol to explore environmental and genetic factors involved in titan cell formation. The genetic determinants of titan cell formation have been investigated through a genotype-phenotype correlation study in H99-derivative laboratory reference strains, through analysis of deletion and complementation in reference strains, and analysis of genetic defects in clinical isolates using whole genome data and complementation.

## Results

### Titan cells generated *in vitro* had similar characteristics to *in vivo* titan cells

While growth in minimal medium using standard growth conditions has no effect on cell size, we identified growth conditions that stimulated the production of enlarged yeast cells and optimized this experimental protocol, referred to as our *in vitro* protocol, using the reference strain H99O (S1 Fig). Observation of these *in vitro*-generated large cells by microscopy shows many characteristics of titan cells including increased cell body size (diameter >10 µm), refractive cell wall, large central vacuole, and peripheral cell cytoplasm distribution, similar to *in vivo* titan cells (Fig 1A). Our *in vitro* protocol proved to be reproducible with H99O reference strains generating titan cells in three different laboratories throughout the world, although some variability in the overall proportion of titan cells generated was observed (Fig 1B). Specifically, the proportion of titan cells was 39.4% [interquartile range (IQR), [34.1-40.7] in Lab 1, 21.0% [12.5-26.8] in Lab 2 and 29.9% [23.1-48.5] in Lab 3. The distribution of yeast cell body size from the *in vitro* protocol varied from 3.7 to 16.3 µm (median 10.2 [8.5-11.5]), whereas *in vivo* it varies from 3.6 to 41.8 µm (median 14.8 [11.2-18.45]) with 84% of the yeasts classified as titan cells (Fig 1C).

**Fig 1.**
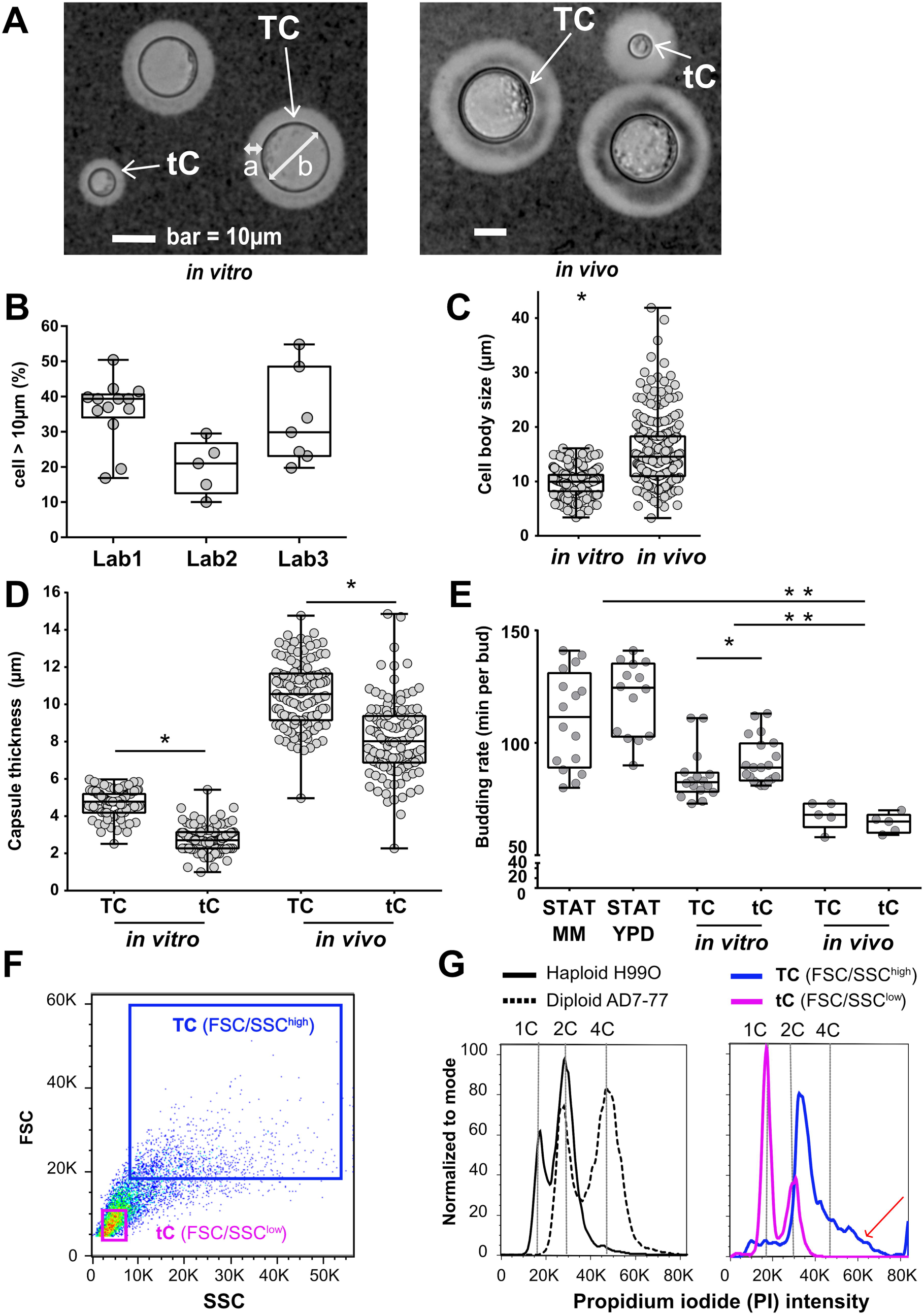
Titan cells generated *in vitro* harbor the typical phenotype of titan cells produced *in vivo*. (**A**) Specific morphology of titan cells (TC, white arrow) *in vitro* (left panel) and *in vivo* (right panel) was observed: enlarged capsule (a), increased cell body size > 10 µm (b), thickened cell wall, large central vacuole, and peripheral cell cytoplasm distribution) while the size of typical cells (tC, white arrow) is <10 µm. (**B**) Titan cells were reproducibly generated in lab 1 using H99O and in two independent laboratories (lab 2 and lab 3) using their local H99O strain. Cell size was measured manually or by using the Icy software on pictures taken in bright field. Each dot represents an independent experiment (median [interquartile range, IQR] are presented); (**C**) Cell body size is increased *in vivo* compared to *in vitro*. Dots represent individual cells, and boxes median and IQR for 230 cells each (*p<0.0001) (**D**) Capsule size measured after India ink staining was significantly larger in titan cells (TC) than in typical cells (tC) both *in vitro* and *in vivo*, in general, the capsule was larger *in vivo* independently of the cell size (p<0.0001). Dots represent individual cells, and boxes median and IQR for 250 cells each (*p<0.001); (**E**). The budding rate of titan cells was lower compared to that of typical cells upon incubation in minimal medium (MM) after titan cells generation *in vitro* *(p=0.018) but not *in vivo*. *In vivo*, budding rate of titan cells (TC) and typical cells (tC) were equivalent and increased as compared to *in vitro* titan cells and typical cells and controls (**p<0.001).(**F**) The yeasts recovered at step 4 of the protocol and analyzed by dot plots (FSC/SSC) using flow cytometry included two populations FSC/SSC^high^ and FSC/SSC^low^ representing titan cells (TC) and typical cells (tC), respectively; (**G**) DNA content analysis after propidium iodide (PI) staining showed that the titan cells (FSC/SSC^high^) population harbored increased PI fluorescence intensity from 2C to >4C (red arrow) as compared to the diploid control (AD7-77 cultured in Sabouraud agar) while the typical cells (FSC/SSC^low^) population harbored a PI intensity comparable to the haploid control (H99O cultured in Sabouraud agar).

Titan cells differ from typical cells in various characteristics including capsule size budding rate, DNA content, cell wall and capsule structure, and the extent of melanization [12,13,20,22]. Comparison of *in vitro* titan cells (TC) to typical cells (tC) showed a significant increase in capsule size (median 4.8 µm in titan cells vs 2.7 µm in typical cells, p<0.001) similar to that observed *in vivo* (median 10.5 µm in titan cells vs 8.0 in typical cells, p<0.001, Fig 1D). The capsule thickness of *in vivo* titan cells was increased compared to the *in vitro* titan cells. The budding rate of *in vitro* titan cells was also significantly increased (median 82.5 m per bud) compared to typical cells (median 89.0 m per bud) (p=0.018), whereas it was similar for titan and typical cells produced *in vivo.* Interestingly, overall budding rate was faster *in vivo* than *in vitro* cells (68 and 65 m per bud, p<0.001)). The budding rate of both in vivo and in vitro titan cells and typical cells were faster than cells grown in stationary phase (111.5 and 124.5 m per bud in minimal medium (MM) and YPD, respectively) (Fig 1E, S1 Movie, S2 Movie). To analyze DNA content, yeasts obtained at the end of the *in vitro* protocol were stained with propidium iodide (PI) and DNA content of titan (TC, FSC/SSC^high^) and typical (tC, FSC/SSC^low^) cells was compared to haploid (H99O) and diploid (AD7-77) strains grown in Sabouraud medium. DNA content was higher in titan (FSC/SSC^high^) than typical (FSC/SSS^low^) cells (Fig 1F), with an increase in the proportion of polyploid cells in the titan cell population as observed by a PI fluorescence greater than the diploid control strain (red arrow, Fig 1G). In contrast, the typical cells had the same PI fluorescence pattern as the haploid H99O cells. Similarly, a reverse gating strategy based on PI intensity shows yeasts with the highest PI intensity were large titan cells (red arrows, S2 Fig).

Calcofluor white (CFW) staining was used to analyze cell wall chitin content. After multispectral imaging flow cytometry – gating on the titan and typical cell populations under both *in vitro* and *in vivo* conditions (Fig 2A-B) – CFW fluorescence intensity (Fig 2C-D) showed significantly increased fluorescence of titan cells compared to typical cells *in vitro* (322539 ± 3072 vs 123062 ± 20727, p<0.0001, Fig 2C) and *in vivo* (144909 ± 38487 vs 27622 ± 7412, p<0.0001, Fig 2D). Cell sorting based on CFW staining and fluorescence microscopy allowed us to validate that cells exhibiting the higher CFW intensity were titan cells (FSC/SSC^high^, S3B Fig). Measurements of chitin content using fluorescence microscopy also showed significant increases in titan cells compared to typical cells (fluorescence intensity/pixel/cell 87.9 [71.7-107.7] vs 66.5 [51.7-79.4], respectively, p<0.0001, S4A Fig). Chitin levels were also assessed by N-acetylglucosamine (Gluc-NAc) content. Gluc-NAc levels were higher in titan cells (156.3 mM/g [153.7-203.7]) than typical cells (97.7 [87.2-119.3]), (p<0.001, FigS4B). Furthermore, titan cells exhibited pronounced melanization (S4C Fig), as measured by blackness on the pictures (Fig S4D), compared to typical cells (S5D Fig), with a median of the (max – mean grey intensity per pixel) of 20065 [18785-21887] in titan vs 13067 [9660-15998] in typical cells (p<0.0001).

**Fig 2.**
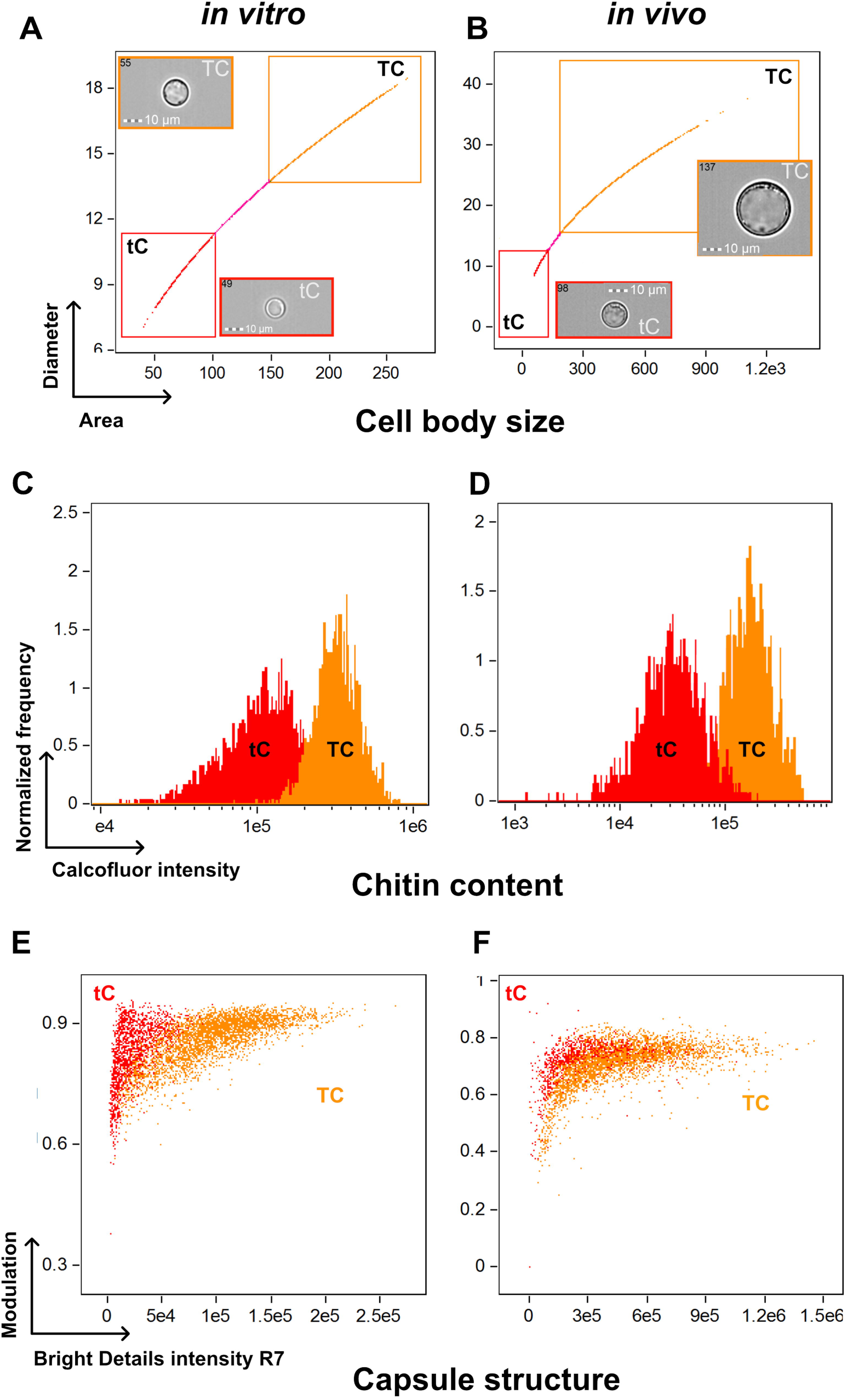
Titan cells harbor an increased chitin content and a specific capsule structure *in vitro* and *in vivo* using multispectral imaging flow cytometry. (**A**) Titan cells (TC) and typical cells (tC) were selected in the corresponding gates based on the Area/Diameter dot plot. **(B)** Titan cells *in vivo* are bigger than that produced *in vitro*. The chitin content based on calcofluor white (CFW) fluorescence intensity showed significantly increased fluorescence of titan cells compared to typical cells *in vitro* (**C**) and *in vivo* (**D**). Based on the fluorescence pattern of the 2D10 anti-capsular monoclonal antibody, the algorithms “modulation” and “bright details intensity R7” allowed to discriminate the capsule structure of titan cells and typical cells with almost no overlap between both population *in vitro* (**E**) and *in vivo* (**F**).

Finally, capsule structure was also investigated based on the binding pattern of monoclonal antibodies specific for capsular polysaccharides [28] using multispectral imaging flow cytometry (Fig 2E-F, S5A Fig) and immunofluorescence (S5B Fig). Based on the fluorescence pattern of the 2D10 antibody [29], the algorithm modulation and bright details intensity R7 allowed us to discriminate the distribution of the capsule staining in titan (TC) and typical (tC) cells and showed with almost no overlap between both population *in vitro* (Fig 2E) and *in vivo* populations (Fig 2F). Variability of staining was observed for E1 (IgG1) and 13F1 (IgM) antibodies (S5A Fig). No pronounced differences between the capsule structures of titan and typical cells was observed visually during immunofluorescence staining (S5B Fig).

### Titan cells develop from older cells and produce typical sized daughter cells

To examine temporal changes in cell size induced by our protocol, we measured yeast cell sizes at 0, 4, 8, 16, 24 and 120 h using automated analysis. This automated analysis correlated with manual size measurements (Interclass correlation=0.99) and titan/typical cell classification (Kappa test=0.81±0.07). The median cell size increased during the first 24 h of incubation, starting at 5.7 µm [5.4-6.0] and increasing to 9.7 µm [8.4-11.0] (Fig 3A). The first titan cells were observed at 8 h with a progressive increase in the proportion of titan cells overtime, reaching a plateau by 24 h (Fig 3B).

**Fig 3.**
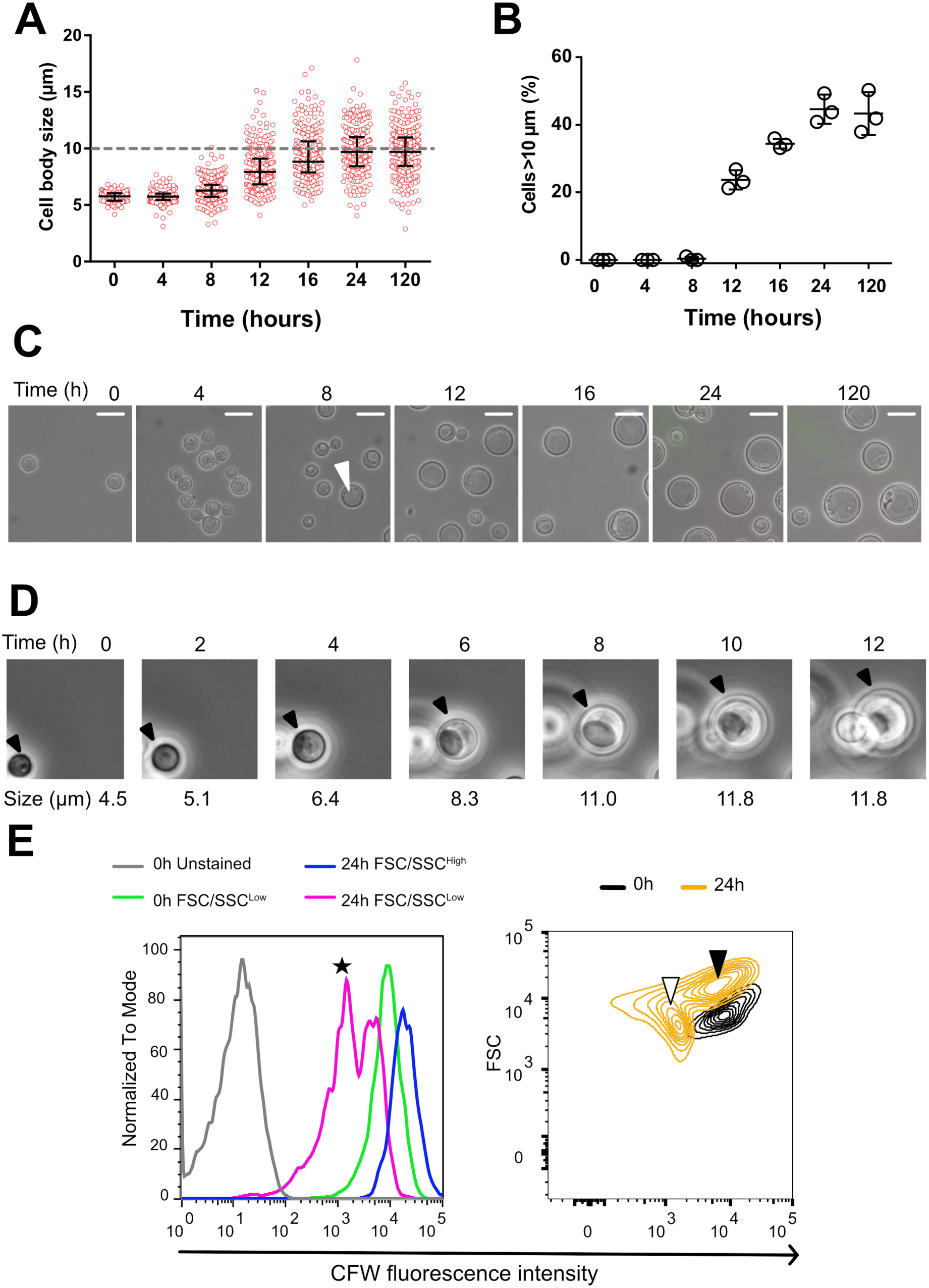
Dynamics of titan cells generation *in vitro*. (**A**) Cell body size was measured from samples of H99O culture (step 3 of the protocol) withdrawn at specific times (H0 to H120) using pictures taken in bright field and measured with the ICY software (mean number of yeasts counted ± SD = 219 ± 67, representative of three experiments). Cell size increased starting at H8 with some cells reaching the threshold of 10 µm (grey dashed line). Each dot represents a single cell and the bars represent median and IQR; (**B**) Titan cells generation started between H8 and H12 and reached a plateau at H24. Each dot represents the proportion of titan cells in the corresponding sample (3 independent experiments); (**C**) Pictures (x400 magnification) taken overtime showing the progressive increase in cell body size and the appearance of a vacuole typical of titan cells in a large cell at H8 (white arrow) (scale bar 10 µm); (**D**) Time lapse imaging of titan cells generation over 12 h showing that titan cells swelled progressively from a small cell and produced daughter cells after having increased their size. (**E**) Cells stained with calcofluor (CFW) at 0.1 µg/mL prior to incubation using our protocol. CFW fluorescent intensity is analyzed by flow cytometry in the initial (H0) and the resulting FSC/SSC^high^ (titan cells) and FSC/SSC^low^ (typical cells) observed at H24. The initial (H0) (green line) and the H24 FSC/SSC^high^ (blue line) populations harbored a high CFW fluorescence suggesting that they are mother cells, with a higher fluorescence for the FSC/SSC^high^, while two populations of high and low (black star) CFW fluorescence intensity were observed for the H24 FSC/SSC^low^ cells (Left panel). The right panel shows the size (FSC) and CFW fluorescence intensity of the yeast populations at H0 (black content lines) and H24 (yellow content lines). The initial population (CFW^high^/FSC^low^) evolved in two populations, one corresponding to daughter cells (typical cells, CFW^low^/FSC^low^, white arrow), and the other one corresponding to titan cells (CFW^high^/FSC^high^, black arrow).

Temporal changes in cell morphology were determined using light microscopy and live cell imaging. Cells with the large vacuole characteristic of titan cells appeared between 4 and 8 h (Fig 3C, white arrow). Live cell imaging over 12 h (Fig 3D, S3 Movie) showed that: (i) titan cells swelled from the progenitor typical sized cells and (ii) titan cells divided to produce typical sized daughter cells. CFW staining is known to transfer only partially to daughter cells upon division resulting in lower fluorescence in daughter cells while remaining at a high level in mother cells [4, 30]. Pulsed CFW staining was used to further monitor the ancestry of titan and typical cell populations over time using flow cytometry (Fig 3E). The initial CFW stained population (0 h) consisted of typical cells (FSC^low^) with high CFW fluorescence intensity (black density lines). At 24 h, two populations were observed. The titan cell population (FSC^high^) had high CFW fluorescence intensity (black arrow), indicating these cells were generated by the swelling of typical sized cells in the original culture. The second population consisted of typical cells (white arrow, FSC^low^) with low calcofluor fluorescence, consistent with newly formed daughter cells (Fig 3E, right panel). Combined, these data show that the titan cells derived from the initial inoculated cells and daughter cells are typical sized.

### Generation of titan cells *in vitro* was influenced by environmental conditions

We tested several parameters affecting steps 2 and 3 of our protocol described in S1 Fig and identified parameters that significantly influence titan cell generation, as measured by cell size distribution and proportion of titan cells. The first parameter we tested was the growth medium and transition between different growth media (Fig 4A). Initial culture in YPD (step 2) then transfer to MM (step 3) resulted in the highest median cell size at 9.1 µm [6.9-11.1]. Initial culture MM followed by transfer to MM produced fewer titan cells, 25.5% (118/463) vs 39.5% (182/461), although the titan cells tended to be larger with cell body diameters over 20 µm. Light exposure during step 3 also increased both median cell size and proportion of titan cells (Fig 4B). In the light, median cell size was 9.4 µm [7.3-11.4] vs 8.4 µm [7.1-9.9] in the dark, with a titan cell proportion of 41.6% (983/2362) vs 24.3% (1072/4399), respectively (Fig 4B). Incubation temperature at 30°C at step 3 increased cell size distribution compared to 37°C (9.4 µm [7.3-11.4] vs 7.0 µm [6.2-8.2]) as well as the proportion of titan cells (41.6% (983/2362) vs 8.1% (297/3652)) (Fig 4C). The pH of the minimal medium at step 3 also influenced cell size (Fig 4D), with pH=5.5 producing significantly larger cells and proportion of titan cells (9.1 µm [6.9-11.2] and 38.6% titan cells), compared to either lower pH (pH=4: median 5.1 µm [4.4-5.8] (0%)) or higher pH (pH=7: 8.2 µm [7.2-9.4] (16.4%), or pH=8.5: 6.9µm [5.9-7.9] (0.7%)) (Fig 4D). Finally, hypoxia at step 3 also increased median cell size compared to normoxia (7.5 µm [5.9-9.7]), with chemically induced hypoxia yielding higher median cell sizes compared to physically induced hypoxia] (10.1 µm [7.8-12.5] vs 8.9 µm [7.3-10.9, p<0.0001) (Fig 4E). The proportion of titan cells in normoxia (14.5% (732/5050) was lower than in chemically induced hypoxia or physically induced hypoxia (63.0% (732/1161) and 38.6% (1264/3004), respectively) (p<0.0001).

**Fig 4.**
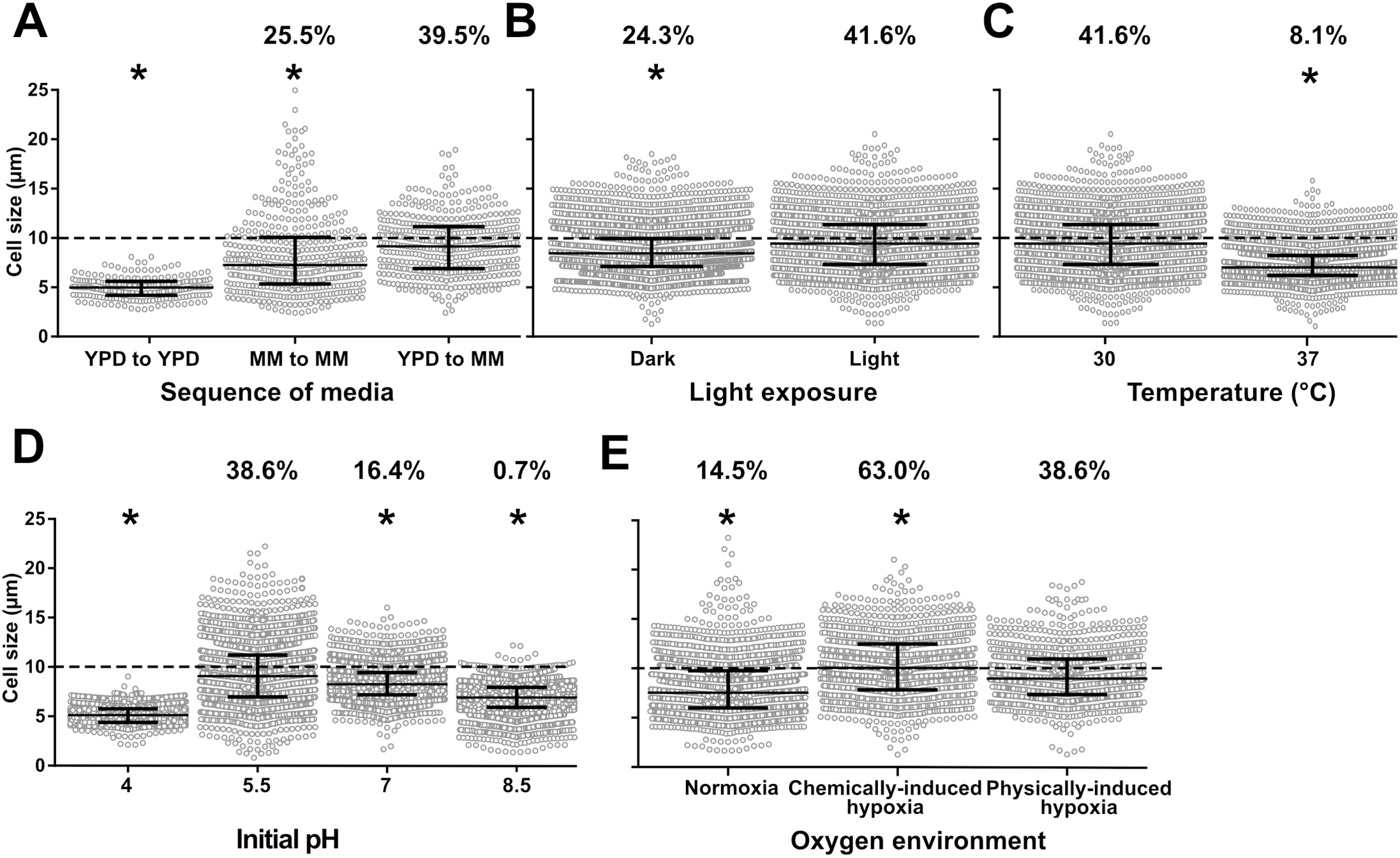
Titan cells generation *in vitro* is impacted by various environmental conditions. (**A**) The sequence of media used at steps 2 and 3 of the protocol was crucial for titan cells generation: yeasts cultured in Yeast Peptone Dextrose (YPD) and transferred to minimal medium (MM) produced significantly more titan cells (cells >10µm, dotted grey line) than yeasts cultured in MM or YPD and transferred in MM or YPD, respectively; (**B**) Exposure to d light at step 3 had a positive impact on titan cells generation compared to incubation in the dark; (**C**) Raising the incubation temperature to 37°C at step 3 decreased titan cells formation compared to 30°C; (**D**) Modification of the initial pH of the MM used at step 3 modifies titan cells formation with pH 5.5 being optimal while a more acidic (pH=4), a neutral (pH=7) or an alkaline (pH=8.5) pH inhibited titan cells formation; (**E**) The impact of hypoxia was tested by physical (closed cap) and by chemical (COCl2 in MM at 1 nM) method and compared to normoxia (21% oxygen). Physically- and chemically-induced hypoxia enhances the production of titan cells compared to normoxia with a higher proportion of titan cells in chemically-compared to physically-induced hypoxia. All experiments were performed in triplicate and pooled (mean cell counted ± SD =2305±1438). Median and IQR are shown in black for each condition (* p<0.0001 vs reference condition). The percentages above each condition represents the % of titan cells observed.

### Generation of titan cells *in vitro* is influenced by host derived cues

We then tested hosts factors that could interact *in vivo* with yeast cells in the lung such as anticapsular antibodies, serum and phosphatidylcholine. Both serum and phosphatidylcholine have already been implicated in titan cell formation [13,18,31]. Co-incubation at step 3 with monoclonal antibodies that bind to different epitopes of the capsule inhibited titan cell generation, with a decreased cell size of 7.2 µm [6.1-8.3] for E1 mAb, and 6.8 µm [5.9-7.7] for 18B7 mAb compared to the untreated control (8.9 µm [7.1-10.6]) (Fig 5A), and a significantly smaller proportion of titan cells (3.2% (44/1360) with 18B7, 5.3% (67/1273) with E1 compared to 33.1% (327/987) for the untreated control). The addition of fetal calf serum (FCS) significantly decreased median cell size (7.3 µm [6.4-8.1] vs (9.1 µm [7.1-11.1], Fig 5B) and the proportion of titan cells (2.5% (99/3911) vs 38.2% (1234/3228), p<0.0001) compared to control, as did the addition of phosphatidylcholine (PC) (8.0 µm [7.0-9.0] vs 9.0 µm [7.1-11.2] for the median cell size, (Fig 5C) and the proportion of titan cells (14.7% (344/2340) vs 38.2% (1077/2820), p<0.0001).

**Fig 5.**
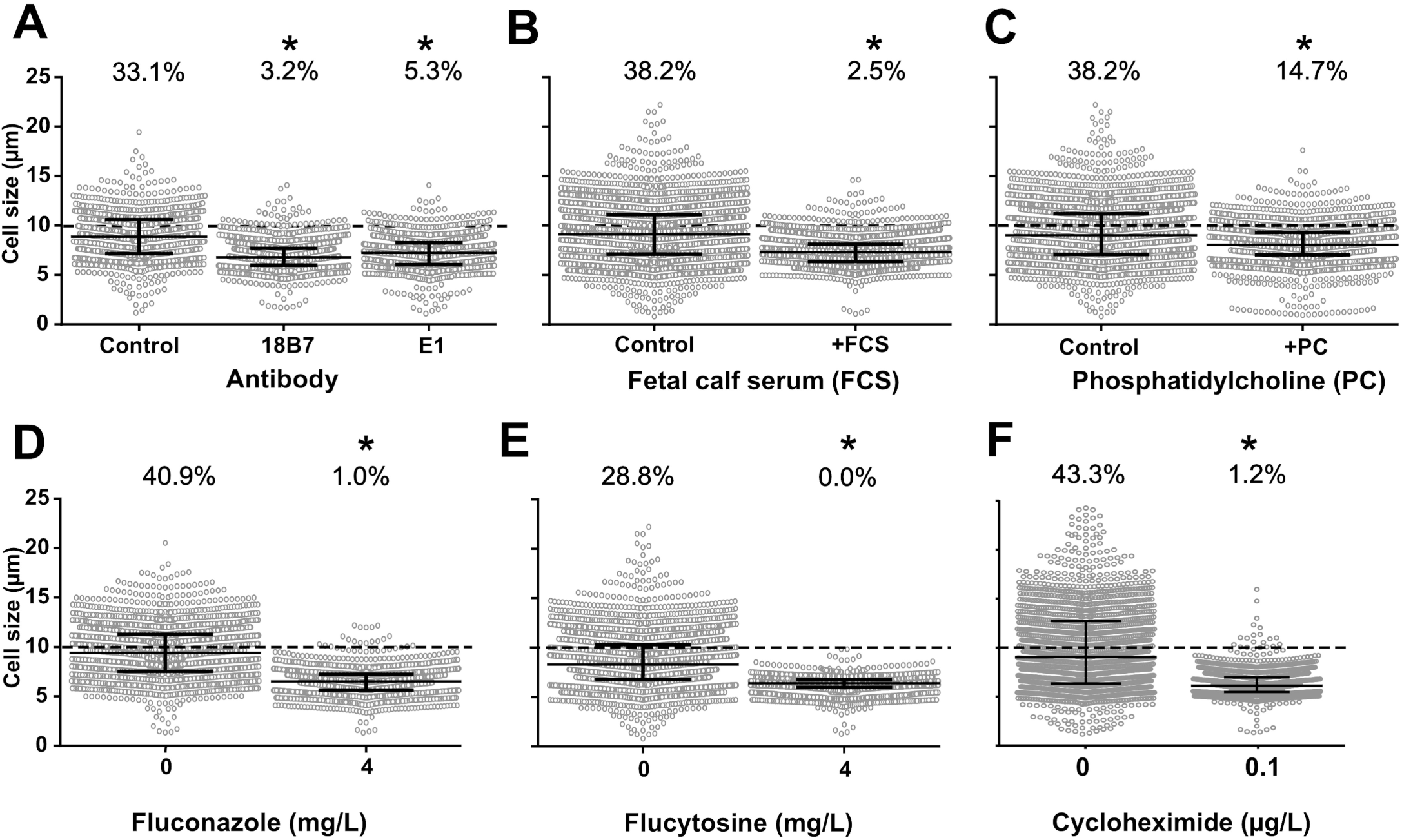
Titan cells generation *in vitro* is influenced *in vitro* by host derived cues and ergosterol, protein and RNA inhibitors. (**A**) Monoclonal anti-cryptococcal capsular polysaccharide antibodies E1 and 18B7 (at 166 µg/mL in MM) significantly decreased cell size, as fetal calf serum (FCS, 5%) (**B**) and phosphatidylcholine at 5mM (**C**) did. (**D**) Fluconazole; (**E**) flucytosine at concentration below the MIC (4 mg/L) and (**F**) cycloheximide at 0.1µg/mL drastically impaired titan cells formation with almost no titan cells produced upon drug exposure.

To understand if an alteration in yeast metabolism induced by ergosterol, protein or nucleic acids inhibition affected titan cell formation, we tested the effect of co-incubation of fluconazole (inhibitor of ergosterol synthesis) and flucytosine (inhibitor of nucleic acids formation and transcription) and cycloheximide (translation inhibitor) at step 3. Fluconazole (FLC) exposure resulted in significantly smaller median cell sizes compared to the drug-free control (7.1 µm [6.2-8.3]) at 1 mg/L, 6.8 µm [5.9-7.8] at 2 mg/L, and 6.5 µm [5.6-7.2] at 4 mg/L vs 9.4 µm [7.5-11.3] in the control, Fig 5G) and a significantly smaller proportion of titan cells (5.9% (124/2073) at 1 mg/L, 2.9% (55/1919) at 2 mg/L and 1.0% (19/1877) at 4 mg/L vs 40.9% (877/2146) in the control, p<0.0001, Fig 5D). Flucytosine exposure significantly decreased the mean yeast cell size at all concentrations tested (6.5 µm [5.9-6.9] at 1 mg/L, 6.5 µm [6.1-6.9] at 2.5 mg/L, and 6.4 µm [5.9-6.8] at 5 mg/L compared to control (8.3 µm [6.8-10.3], Fig 5E) with no titan cells observed upon flucytosine exposure. Cycloheximide exposure at 0.1 µg/mL also significantly decreased the mean cell size from 9.0 µm [6.3-12.7] to 6.1 µm [5.5-6.9] (p<0.0001) and the proportion of titan cells from 43.3% (797/1840) to 0.1% (20/1663) (Fig 5F). Of note, the viability of the cells recovered at step 4 after 5 d of drug exposure was unchanged for fluconazole but reduced for flucytosine and cycloheximide (p<0.0001 compared to unexposed, S6 Fig).

We also tested if iterative subcultures with or without the presence of active molecules (CFW or fluconazole) affected titan cell formation, assuming that the cell wall and the global metabolism of the sub-cultured progeny would be impaired in the presence of high concentrations of the cell wall toxic drug (CFW) or fluconazole, respectively. We analyzed the impact of repeated sub-culture of the cells prior to step 1 on titan cell production (S7 Fig). Sub-culture on Sabouraud agar eight times (8 Sub) spanning a one-month period significantly decreased the median cell size (8.60 µm [7.02-10.07]) compared to the initial culture (0 Sub) (9.17 [6.99-10.90]) (p<0.0001). Addition of CFW to induce cell wall stress during sub-culture (8Sub+CFW) significantly decreased the median cell size (8.03 µm [6.82-9.46]) compared to the 8Sub control. Thus, iterative exposure to fluconazole during sub-culture significantly increased the median yeast cell size to 10.15 µm [8.04-13.23] (8Sub+FLC) compared to the 8Sub control (p<0.001, S7 Fig).

### Generation of titan cells *in vitro* is influenced by quorum sensing molecules

Previous studies in a murine pulmonary infection model showed that inoculum concentration can impact titan cell production [12, 13]. To explore this phenomenon further, we examined cell size changes in response to different initial concentrations of cells at step 3 (Fig 6A). Initial cell concentrations significantly impacted the median cell size of the yeast population (p<0.0001), with the highest median cell size observed at 10^6^ cells/mL (9.2 [7.3-11.1]) compared to 10^5^ cells/mL (6.3 [5.1-8.3]), 10^4^ cells/mL (6.2 [5.1-7.9]) and 10^7^ cells/mL (5.9 [5.2-6.5]) (Fig 6A). Similarly, the proportion of titan cells was significantly higher at 10^6^ cells/mL (37.6% (896/2382) compared to 10.3% (279/2716) at 10^4^ cells/mL, 14.3% (346/2420) at 10^5^ cells/mL and 0% at 10^7^ cells/mL (0/2177), p<0.0001. Previous study reported that pantothenic acid (PA vitamin B5) is involved in quorum sensing and growth rate in *C. neoformans* [32]. The addition of PA had no effect on median cell size (8.35 µm [6.9-10.2] and 8.3 µm [6.4-10.8], p=0.8011, Fig 6B), but significantly increased the proportion of titan cells (37.9% (1435/3785) vs 26.9% (983/3650)). In specifically implemented experimental settings, the proportion of titan cells was influenced by the concentration of PA with a significant increase in titan cells at 0.125 µM (56.5 [50.6-61.1] and 12.5 µM (47.6 [35.6-50.3]) (Fig 6D). In parallel, analysis of the growth curves of the yeast showed a significant increase in the doubling time (slope) at ≥0.125 µM of PA (Fig 6E), suggesting a lack of correlation between titan cell formation and growth rate because titan cell formation was completely inhibited at 1250 µM of PA while the doubling time increased.

**Fig 6.**
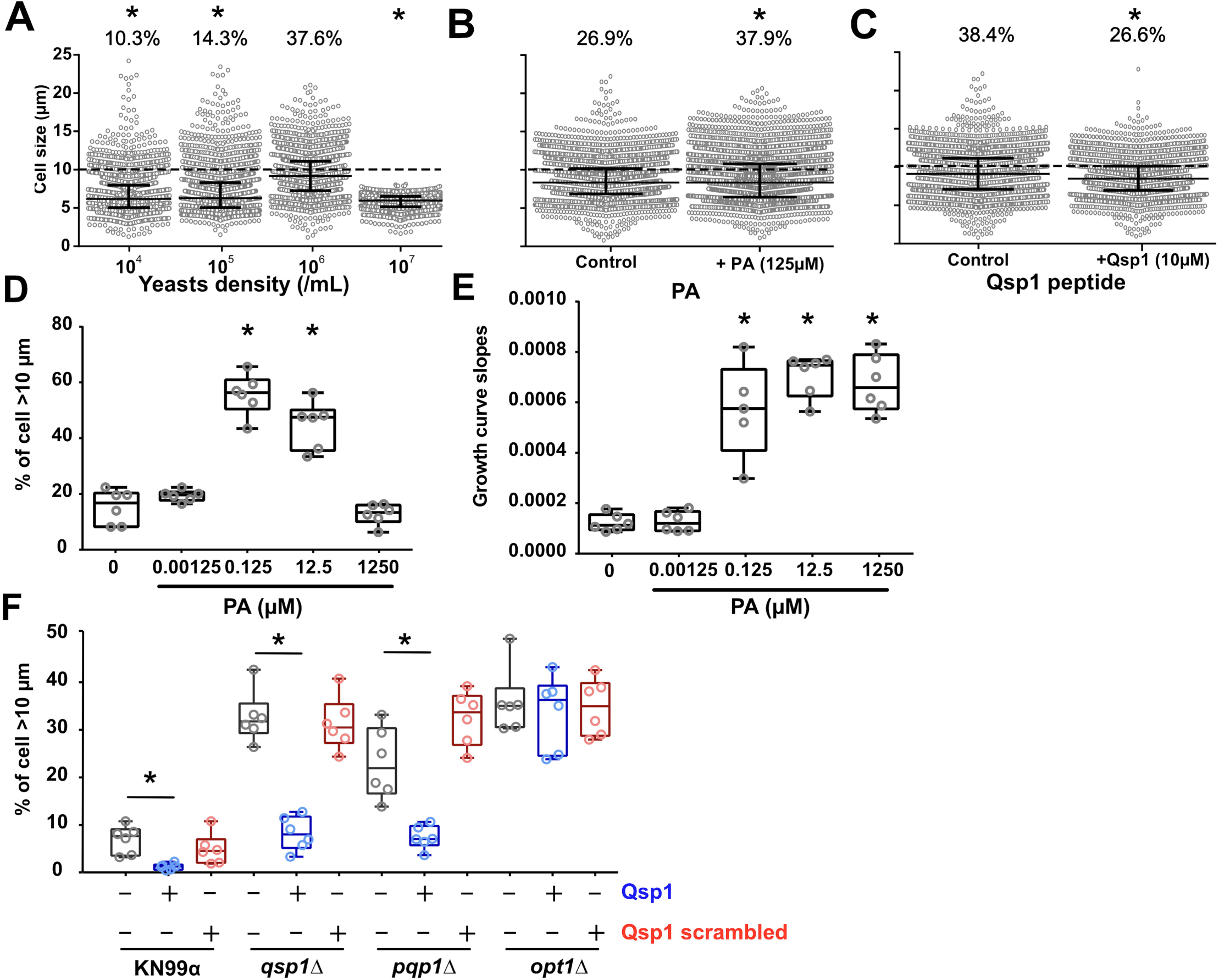
Generation of titan cells *in vitro* is influenced by cell concentration and quorum sensing molecules. (**A**) The cell concentration at onset of step 2 significantly modified titan cells generation with a maximal titan cells formation using an initial concentration of 10^6^ cells/mL, and an abolition of titan cells formation at 10^7^ cells/mL; We then tested several factors at step 3 by adding (**B**) Pantothenic acid (PA) at 125 µM which had no impact on cell size distribution but significantly increased the proportion of titan cells produced; (**C**) quorum sensing peptide 1 (Qsp1p, 10µM) which significantly decreased cell size distribution and the proportion of titan cells produced; (**D**) The proportion of titan cells generated is influenced by the concentration of PA with a significant increase of titan cells at 0.125 µM and 12.5 µM (* p<0.001). (**E**) Growth curves were measured continuously during titan cells generation and showed a significant increase of the doubling time (slope) from 0.125 µM of PA on. titan cells formation and growth rate is not correlated since titan cells formation is inhibited at 1250 µM of PA and the doubling time increased. (**F**) Qsp1 acts as a repressor of titan cells formation as the addition of Qsp1 peptide inhibited titan cells formation (*p<0.001). A control using scrambled peptide showed no effect on titan cells formation. The *qsp1Δ* and *pqp1Δ* complemented by addition of Qsp1p showed an increase in titan cells formation. No effect of the Qsp1p was observed on *opt1Δ*, the deletion mutant of the Qsp1 transporter Opt1. A specific titan cells inducing conditions was implemented for D, E, and F to allow induction of titan cells in a 100-well plate and continuous measurements of growth curves using the Bioscreen apparatus.

Recent studies in *C. neoformans* also implicate the role of the small Qsp1 peptide in quorum sensing [33]. Addition of Qsp1 peptide significantly decreased median cell size from 9.1 µm [7.1-11.2] to 8.5 µm [6.9-10.1] (Fig 6C) and titan cell proportion from 38.4% (1075/2798) to 26.6% (915/3439), p<0.0001. Addition of Qsp1 peptide inhibited the formation of titan cells in H99O (Fig 6C) and KN99α (Fig 6F). In the *qsp1Δ*, *pqp1Δ* and *opt1Δ* deletion mutants that cannot produce or import a functional Qsp1 peptide [33], titan cell generation was increased compared to KN99α, confirming the negative regulation of Qsp1 peptide in titan cell formation (Fig 6F). When *qsp1Δ*, *pqp1Δ* were complemented with Qsp1 but not with scrambled Qsp1 peptides, titan cell formation was similar (increased titan cell formation) to that of the mutant alone. The complementation of the *opt1Δ* deletion mutant with Qsp1 or scrambled Qsp1 did not rescue the parental phenotype suggesting that the import of Qsp1 is crucial for its action on the yeast cells (Fig 6F).

### *In vitro* titan cell generation is dependent upon H99 genetic background and requires functional *LMP1, SGF29* and *SREBP* genes

Previous whole genome sequencing studies identified single-nucleotide polymorphisms (SNPs) and insertions/deletions (indels) between H99-derived strains recovered from various laboratories (Table 1) [34]. To determine whether any of these SNPs or indels affected titan cell generation, we tested the H99S, H99W, H99 CMO18, H99L, KN99α strains. H99O produced significantly more titan cells than the other H99-derived strains, p<0.0001 (Fig 7A, S8A-S9 Fig). These H99 derivative strains were also tested for titan cell formation in the lungs of infected mice (S8A-S9 Fig). As with *in vitro* titan cell production, all the H99 derivative strains showed lower levels of titan cell formation *in vivo* when compared to H99O (p<0.0001), with the exception of KN99α that had equivalent titan cell production to H99O (Fig 7A, S8A-S9 Fig).

**Fig 7.**
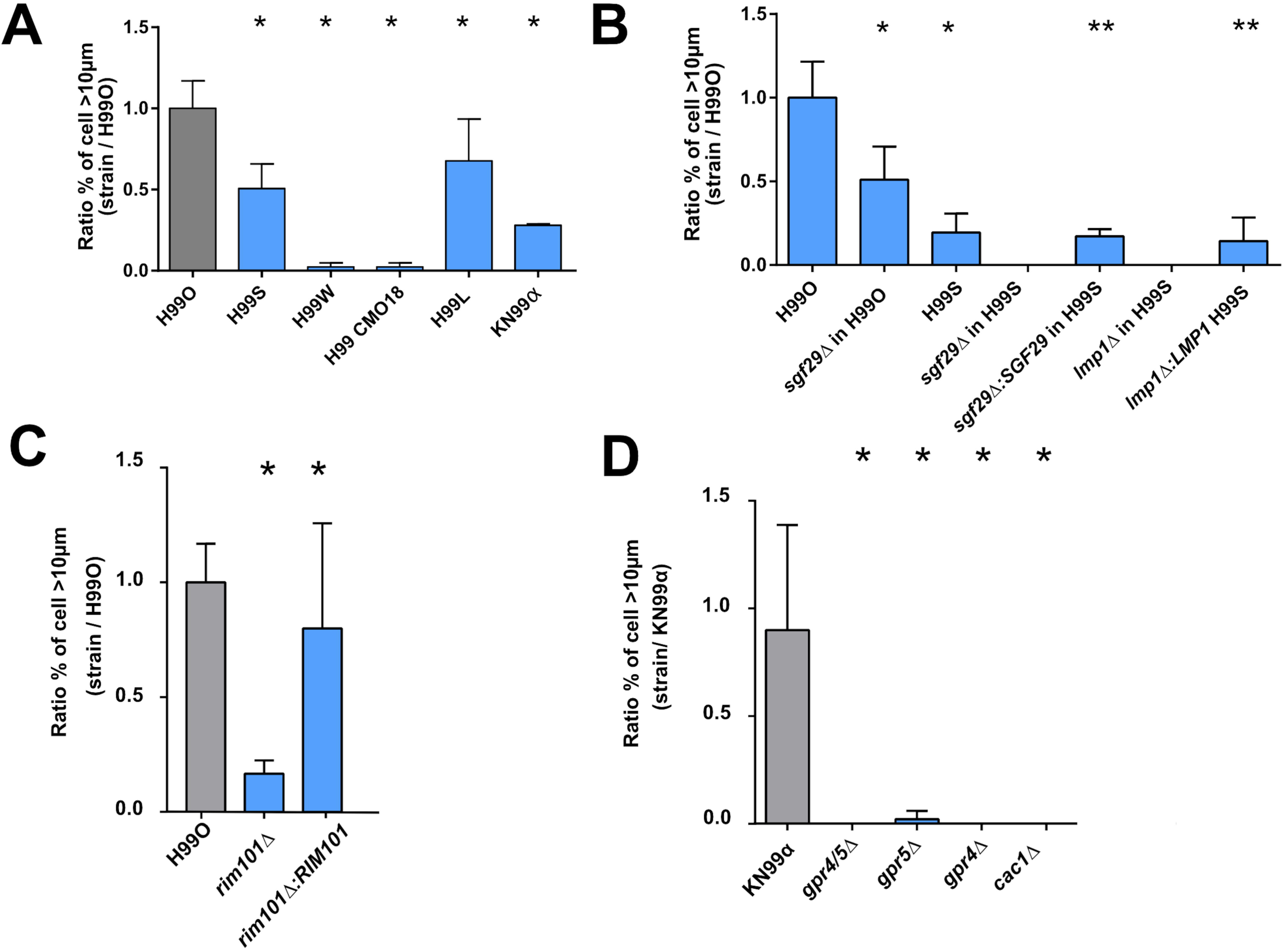
Titan cells generation is dependent on various genes and requires signaling through the Gpr/PKA/Rim101 pathway *in vitro*. (**A**) The different H99 strains harbored variable abilities to produce titan cells compared to H99O (grey bar) with high titan cells producer (H99O, S, L) and low titan cells producer (KN99α, H99W, H99 CMO18) *in vitro*. (**B)** *Sgf29Δ* and *lmp1Δ* deletion mutants show a decrease in titan cells generation in various H99 backgrounds *in vitro* compared to H99O. Complementation in strains *lmp1Δ:LMP1* and *sgf29Δ:SGF29* in H99S background restored the phenotype of H99S. Rim101 (**C**) and *GPR4* and *GPR5* and *CAC1* **(D)** are required for titan cells generation *in vitro* in H99 and KN99α. The ratio to the value obtained for H99O used as a calibrator in each experiment was calculated for each strain. Bar represent mean ± SD (mean cell counted=600). Khi2 test was performed to compare the experimental conditions to H99O, they were performed in triplicates and pooled (*p<0.0001, **p<0.0001, when the comparison was done with the parental strain H99S.)

**Table 1:** Genes affected by SNPs or Indels in the different H99 strains.

| Strains | Genes affected | Gene function | Genetic event |
| --- | --- | --- | --- |
| H99O | *CNAG_07634* | hypothetical | Deletion |
|  | *CNAG_04078* | hypothetical | Insertion |
|  | *CNAG_06456* | hypothetical | Insertion |
| H99S | *CNAG_07595* | hypothetical | Deletion/frameshift |
|  | *CNAG_12447* | miscRNA | Insertion |
|  | *CNAG_12900* | miscRNA | Insertion |
| H99L | *CNAG_06392, SGF29* | transcription factor (binds H3K4me2/3 and recruits histone deacetylation) | Deletion |
| KN99α | *CNAG_06392, SGF29* | transcription factor (binds H3K4me2/3 and recruits histone deacetylation) | Deletion |
| H99W | *CNAG_06765, LMP1* | Hypothetical (involved in mating virulence and melanization) | Deletion/frameshift |
| H99 CMO18 | *CNAG_06765, LMP1* | Hypothetical (involved in mating virulence and melanization) | Deletion/frameshift |

Two genes, *LMP1* and *SGF29*, are dramatically affected by SNPs/indels in the H99 derivatives; *LMP1* has a frameshift deletion (H99W and H99 CMO18) and *SGF29* is deleted (KN99α and H99L) [35]. To determine if these genes are involved in titan cell production, we analyzed *lmp1Δ* and *sgf29Δ* deletion mutants for *in vitro* and *in vivo* titan cell formation (Fig 7 and S8-S9 Fig, respectively). In vitro, the *sgf29Δ* mutant in the H99O background had half the titan cell formation of the H99O wild-type strain [8.1% (49/600) to 4.2% (25/600), p<0.0001]. The *sgf29Δ* mutant in the hypervirulent H99S also manifested no titan cells generation, as did an *lmp1Δ* mutant in this background. Complementation of *LMP1* and *SGF29* in this H99S mutant restored titan cells generation to that found in the parental strain; 1.2% (13/935) and 1.4% (7/600), respectively vs H99S 1.6% (25/1540) (Fig 7B). Importantly, the same trend was observed for *in vivo* titan cell formation*. In vivo*, the *lmp1Δ* H99S mutant produced only 3.5% (21/600) titan cells compared to 14% for H99S (84/600), p<0.0001, and this decrease in titan cell production was restored in the *lmp1Δ:LMP1* H99S strains (9.5% (57/600)) (S8B Fig). The *sgf29Δ* mutant in the H99O background reduced titan cell formation *in vivo* from 18.8% (113/600) to 9.3% (37/400) (p<0.0001). In H99S, complementation of Sgf29 *(sgf29Δ:SGF29*) in H99S restored titan cell generation to wild-type H99S levels from 5% (30/600) to 19.8% (237/1200) (Fig 7B, S8B Fig) (p<0.0001).

SREBP is a gene involved in response to hypoxia, so we tested titan cell formation in the *sre1Δ* mutant. The proportion of titan cells was significantly decreased in the *sre1Δ* mutant at 5.1% (53/920) compared to KN99α [14% (337/2358)] (p<0.0001) (S10 Fig).

### *In vitro* titan cell generation requires signaling through the Gpr/PKA/Rim101 pathway and is dependent on negative regulators

The signal transduction pathway Gpr/PKA/Rim101 regulates titan cell formation *in vivo* [18]. Briefly, the G-protein coupled receptor 5 and Ste3a pheromone receptor signal through Gpa1 to trigger the cAMP/PKA signaling cascade, ultimately activating the Rim101 transcription factor. This pathway regulates virulence factors such as capsule or melanin [36, 37].

To determine if this same pathway was critical for titan cell generation *in vitro*, we examined cell enlargement in the *gpr4Δ*, *gpr5Δ*, *gpr4Δ*/*gpr5Δ, rim101*Δ, and *cac1Δ* mutants and their complemented strains in both the H99 and KN99α genetic backgrounds (Fig 7C-7D, S9 Fig). In the H99O genetic background, Rim101 function was similar to that observed *in vivo*, with little titan cell formation in the *rim101Δ* mutant (1.9% (51/2600)) and full restoration of titan cell production in the complemented strain (9.2% (239/2600) (p<0.0001) (Fig 7C). In KN99α, the *rim101Δ, gpr4Δ/gpr5Δ*, and *cac1Δ* mutants had no titan cell formation, but surprisingly both of the single *gpr4Δ* and *gpr5Δ* mutants also lacked titan cell formation (Fig 7D). This is in contract to *in vivo* where titan cell production was rescued by GPR5 alone [18]. Taken together, these data suggest that signaling through both Gpr4 and Gpr5 via the cAMP/PKA pathway to Rim101 is required for titan cell production *in vitro*.

### *In vitro* titan cell formation is regulated by PKR1 in clinical isolates

To determine whether the Gpr/PKA/Rim101 pathway can impact titan cell formation in clinical isolates, we also screened a total of 56 clinical isolates for their ability to produce titan cells. Two isolates (AD2-06a and AD2-02a) produced a more titan cells relative to H99O (ratio of clinical strain/H99O of 2.6±0.3 and 1.4±0.3, respectively). Three additional isolates (AD4-37a, AD1-95a, AD4-43a) produced fewer titan cells than H99O (ratio=0.4±0.3, 0.2±0.1, 0.1±0.0, respectively). Titan cell production in the other clinical isolates was close to zero (ratio between 0.1 and 0.01% for five, less than 0.01% for six, and no titan cells at all for the remaining 39 isolates) (Fig 8A).

**Fig 8.**
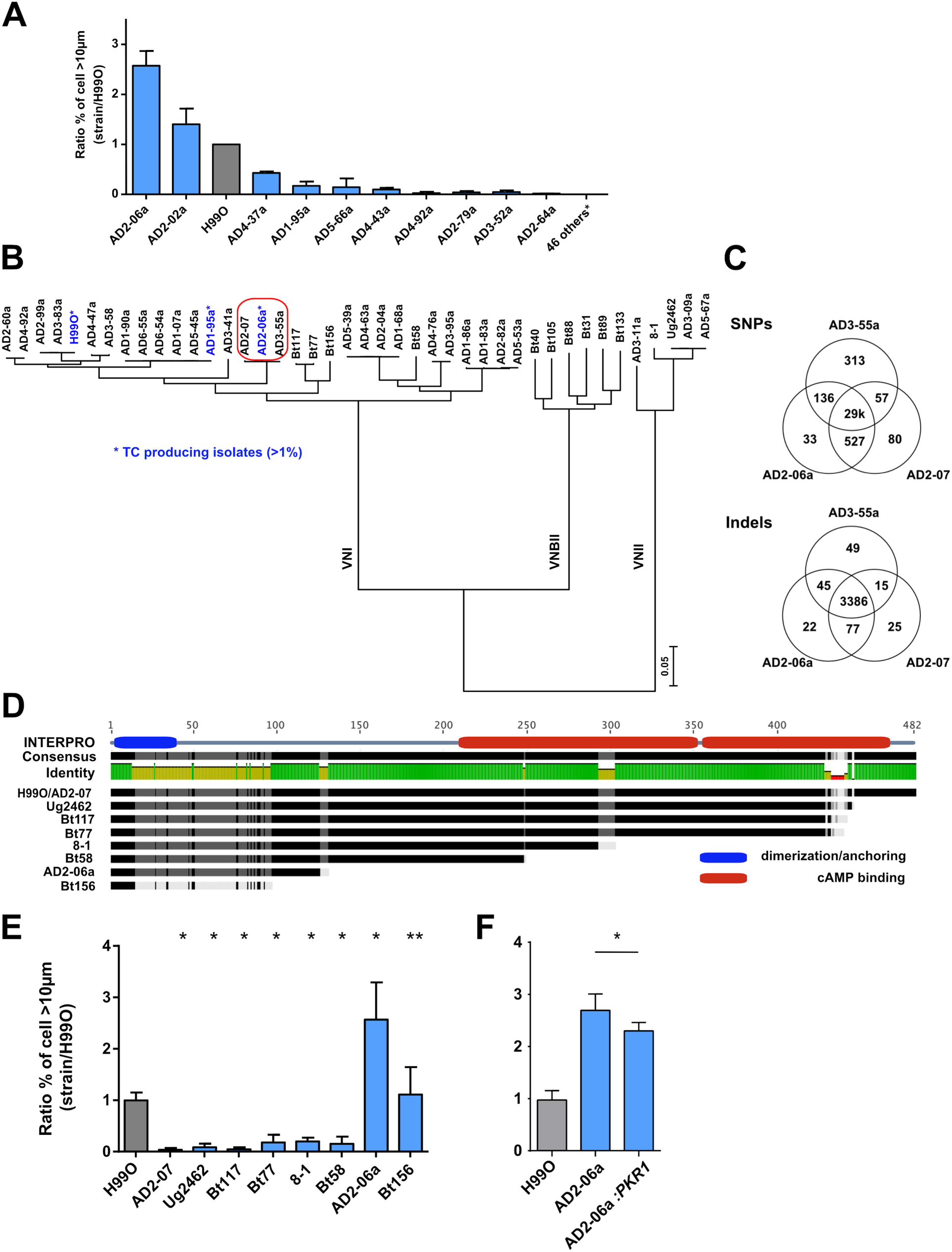
Non-synonymous mutation of *PKR1* enhances titan cells generation based on clinical isolates analysis. (**A**) The screening of 56 *C. neoformans* serotype A MATα french clinical isolates identified isolates AD2-06a (ratio=2.6±0.2) and AD2-02a (ratio=1.4±0.2) as titan cells producers compared to H99O (grey bar); (**B**) Phylogenic tree of the isolates (n=41) for which the whole sequence was available [38, 39] was estimated using RAxML under the GTRCAT model. This shows that AD2-06a, AD2-07 and AD3-55a are phylogenetically close together. (**C**) Venn diagram representing the number of common or specific SNPs and Indel between AD2-06a, AD2-07 and AD3-55a. One of the four AD2-06a specific SNPs is a non-synonymous mutation in the *PKR1* gene (mutation Gly125fs); (**D**) Alignment of Pkr1 protein sequences from selected clinical isolates including H99O and AD2-07 that both harbor a wild type sequence. The Pkr1 domain architecture is formed by a dimerization domain in green (amino acids 2 to 40) and two cAMP binding sites based on INTERPRO model in red (amino acids 219 to 351 and 353 to 473). (**E**) Strains with *PKR1* loss-of-function mutation showed a variable ability to produce titan cells, as compared to H99O (grey bar). The clinical isolate AD2-07, which produces significantly less titan cells was recovered from the CSF of an HIV-positive patient on d 13 of amphotericin B treatment while AD2-06a, which produced the highest proportion of titan cells, was recovered from the initial CSF sample of the same patient. (**F**) Complementation of *PKR1* gene (*AD2-06a:PKR1*) in the naturally deficient strain AD2-06a reduced titan cells generation compared to AD2-06a. The ratio to H99O, used as a calibrator in each experiment, was calculated for each strain (panel A, E, F) and results expressed as mean ±SD. Experiments A, F and G were done in triplicate, B twice (screening). To compare the experimental conditions to H99O, Khi2 analysis was performed (*p<0.0001, **p<0.05). Mean cell counted ± SD = 1165±528.

The complete genome sequence was obtained for 41 of the total 68 screened clinical isolates and a phylogenetic tree of these strains and the H99O reference strain shows high genetic diversity including VNI, VNII, VNBII isolates (Fig 8B, Table 2). Compared to H99O, the high titan cells generating strain AD2-06a harbored 31,229 SNPs and was closely related to AD3-55a (31,171 SNPs) and AD3-41a (28,599 SNPs), which were both unable to produce titan cells. A total of 19 genes, including CNAG_00570 (*PKR1*), were disrupted in AD2-06a and not in AD3-55a and AD3-41a (S1 Table). Of note, no common genetic mutation, insertion or deletion has been observed in the 3 strains that produced a significant (>1%) proportion of titan cells (H99O, AD2-06a, AD1-95a) as compared to all the other examined isolates. Duplication of chromosomal regions were observed in the French sequenced clinical isolates (S2 Table) with AD2-06a harboring a large duplication of chromosome 9 (Chr9, region 465 – 665kb). To assess whether the Chr9 duplicated region could be responsible for titan cell formation, we explored a larger collection of *C. neoformans* strains with complete genome sequence [38] and discovered five additional clinical isolates (Ug2459, CCTP20, FFV14, WM-148, WM-626) harboring partial duplications on chr9 duplication (S11 Fig). We analyzed titan cell formation in these 5 isolates, but only the Ug2459 strain generated titan cells *in vitro* and only at a low proportion (S11 Fig), suggesting that genes located within in the AD2-06a Chr9 duplication are not involved by themselves in titan cell formation.

**Table 2.** Alignment and SNP statistics of sequenced isolates using H99 as a reference.

| Strain | MLST genotype | % of<br>Assembly<br>Covered by<br>aligned reads | Alignment<br>depth (X) | SNPs |
| --- | --- | --- | --- | --- |
| AD3-55a | unique | 100 | 133 | 28,070 |
| AD2-06a | unique | 100 | 129 | 28,202 |
| AD2-07 | unique | 99 | 45 | 27,797 |
| AD3-41a | unique | 100 | 134 | 25,378 |
| AD4-92a | A3/M3 | 100 | 132 | 11,906 |
| AD4-47a | A4/M1 | 100 | 132 | 10,018 |
| AD3-58 | A4/M1 | 100 | 123 | 9,996 |
| AD3-83a | A1/M1 | 100 | 166 | 172 |
| AD2-99a | A1/M1 | 100 | 119 | 180 |
| AD6-55a | related to A1/M1 | 100 | 108 | 12,440 |
| AD6-54a | related to A1/M1 | 100 | 131 | 12,440 |
| AD5-45a | related to A1/M1 | 100 | 142 | 12,417 |
| AD1-95a | related to A1/M1 | 100 | 107 | 12,521 |
| AD1-90a | related to A1/M1 | 100 | 122 | 12,511 |
| AD5-67a | VNII | 100 | 139 | 277,381 |
| AD3-9a | VNII | 100 | 111 | 277,417 |
| AD3-11a | VNII | 100 | 125 | 268,842 |
| AD4-76a | Th | 100 | 112 | 43,281 |
| AD3-95a | Th | 100 | 141 | 43,235 |
| AD5-53a | A5/M5 | 100 | 130 | 42,291 |
| AD2-82a | A5/M5 | 100 | 133 | 42,271 |
| AD1-86a | A5/M5 | 100 | 105 | 43,778 |
| AD5-39a | A4/M4 | 100 | 108 | 43,696 |
| AD4-63a | A4/M4 | 100 | 129 | 43,736 |
| AD2-04a | A4/M4 | 100 | 137 | 43,733 |
| AD1-68a | A4/M4 | 100 | 132 | 43,778 |

AD2-06a was isolated from the initial cerebrospinal fluid (CSF) sample of an HIV-infected patient at baseline (diagnosis of cryptococcosis). Another isolate, AD2-07, was recovered from the CSF of the same patient after 13 d of amphotericin B treatment. Thus, AD2-06a and AD2-07 are closely related with only 137 different SNPs and 40 indels. AD3-55a is in the same clade and also very similar with 370 different SNPs and 64 indels, although recovered from another patient and in another place (Fig 8B-8C). By contrast, these three isolates differ from H99O by 29,000 SNP and 3,386 indels. AD2-07 and AD3-55a were both unable to produce titan cells (Fig 8A, 8B, 8E), allowing a more fine-scale analysis of SNPs linked to titan cell formation in these closely related strains (Fig 8C). Specifically, AD2-07 produced 1.0% (31/3047) titan cells compared to 39.1%, (639/1633) in AD2-06a and 15.2% (304/2001) in H99O (p<0.0001). The median cell size of AD2-07 was significantly decreased compared to AD2-06a and H99O (5.9 µm [5.2-6.6] for AD2-07, 8.5 µm [7.0-13.0] for AD2-06a, and 7.7 µm [6.5-9.2] for H99O, p<0.0001, S12 Fig).

Comparison of the AD2-06a, AD2-07, and AD3-55a genomes identified four genes with loss-of-function mutations in AD2-06a but not in AD2-07 or AD3-55a: CNAG_00570 (PKR1) (Fig 8D), CNAG_07475 (hypothetical protein), CNAG_01240 (hypothetical protein) and CNAG_05335 (hypothetical protein). More precisely, AD2-06a had a frameshift mutation at glycine 124 in the CNAG_00570 (*PKR1*) leading to a truncated protein of 138 amino acids (Fig 8D). Pkr1 is the cAMP-dependent protein kinase regulatory subunit that interacts with Pka to regulate the phosphorylation activity of Pka. To further explore *PKR1* in clinical isolates, we analyzed additional, previously sequenced, clinical isolates (S3 Table) that harbored mutations leading to Pkr1 truncation (Bt156, Bt58, 8-1, Bt77, Bt117, Ug2462) for titan cell formation [39]. Specifically, a frameshift mutation at amino acid 14 introducing a premature stop codon at position 96 for Bt156, as well as stop codons introduced at positions 130 for AD2-06a, 258 for Bt58, 302 for 8-1, 439 for Bt77, 441 for Bt117, and 445 for Ug2462 was observed (Fig 8E, Table 3). We hypothesized that the strains with highly impacted/truncated Pkr1 protein would produce more titan cells, similar to the AD2-06a isolate. AD2-06a and Bt156, the strains with the largest truncation, had high levels of titan cell formation with a ratio of 2.8±1.0 for AD2-06a and 1.3±0.6 for Bt156 compared to H99O (Fig 8E), and titan cell proportions and median cell sizes of 39.1% (639/1633) (median 8.5µm [7.0-13.0]) and 17.9% (469/2614) (median 7.7µm [6.4-9.4]), respectively (S12 Fig). For the other strains, the ratio, proportion of titan cells, and median size were decreased compared to H99O: 0.2±0.0, 3.6% (89/2464) and 6.9 µm [6.0-7.8] for 8-1 strain; 0.2±0.1, 2.9% (110/3834) and 6.5 µm [5.8-7.1] for Bt77; 0.1±0.0 0.9% (27/3081) and 6.4 µm [5.7-7.2] for Bt117; and 0.1±0.1, 1.5% (54/3628) 5.4µm [4.7-6.1] for Ug2462, p<0.0001) (Fig 8E, S12 Fig).

**Table 3:** Mutations of *PKR1*, *CAC1*, *USV101* in specific clinical isolates.

| Gene | Strains | Mutations |
| --- | --- | --- |
| *PKR1* | Ug2462 | n.1333C>T\|p.Arg445*\|1333/1449 |
|  | Bt117 | n.1274_1280dupCTCTCCT\|p.Asn428fs\|1280/1449 |
|  | Bt77 | n.1293dupA\|p.Arg432fs\|1293/1449 |
|  | 8-1 | n.878_911delCCGAGGGGAGCTCGTTTGGGGAGTTAGCGCTGAT\|p.Ser293fs\|911/1449 |
|  | Bt58 | n.742G>T\|p.Glu248*\|742/1449 |
|  | AD2-06a | n.375_376delAG\|p.Gly125fs\|376/1449 |
|  | Bt156 | n.41dupA\|p.Asp14fs\|41/1449 |
| *CAC1* | Bt133 | n.41delA\|p.His14fs\|41/6930 |
|  | 8-1 | n.2_3insA\|p.Met1fs\|2/6930 ; n.80G>A\|p.Trp27*\|80/6930 ; n.81G>A\|p.Trp27*\|81/6930 |
|  | AD3-11a | n.2_3insA\|p.Met1fs\|2/6930 |
|  | AD3-9a | n.2_3insA\|p.Met1fs\|2/6930 |
|  | AD5-67a | n.2_3insA\|p.Met1fs\|2/6930 |
|  | Ug2462 | n.2_3insA\|p.Met1fs\|2/6930 |
| *USV101* | Bt88 | n.842dupC\|p.His282fs |

\*, stop codon introduced by the SNP

To directly test whether the truncated Pkr1 protein impacted titan cell production in strain AD2-06a, the functional KN99α allele of the *PKR1* gene was introduced into the strain (AD2-06a*:PKR1*) and titan cell formation analyzed. Titan cell production was significantly decreased by complementation (p<0.0001) with 44.9% (574/1279) for AD2-06a*:PKR1* as compared to 64.6% (700/1083) for AD2-06a and 23.2% (555/2363) for H99O (Fig 8F).

To further explore the function of *PKR1* in titan cell formation, we also tested the ability of *pkr1Δ* in a KN99α background to generate titan cells compared to KN99α, and found that *pkr1Δ* produced more titan cells with a ratio of 4.9±1.6 compared to the parental strain KN99α (Fig 9). The *pkr1Δ* median cell size (8.1 µm [6.7-9.5]) exceeded that of KN99α (6.4 µm [5.5-7.2]) p<0.0001) with a significant increase in the proportion of titan cells at 28.5% (695/2436) for *pkr1Δ* vs 4.6% (121/2614) for KN99α (p<0.0001, Fig 9A). We also analyzed titan cell production in two additional independent *pkr1Δ* and complemented *pkr1Δ:PKR1* strain in the H99 background. The *pkr1Δ-1 and* complemented *pkr1Δ:PKR1-1* gave ratios of 2.9±1.5 and 1.8±0.7, respectively, while the *pkr1Δ-2 and* complemented *pkr1Δ:PKR1-2* ratio were 1.9±0.8 and 1.4±0.4, respectively, compared to the H99 parental strains (1.0±0.5) (Fig 9B). In both strains, complementation significantly reduced the proportion of titan cells generated (p<0.0001) from 29.8% (685/2493) for *pkr1Δ-1* to 18.8% (593/3217) for *pkr1Δ:PKR1-1*; and from 19.9% (422/2364) for *pkr1Δ-2* to 13.9% (359/2588) for *pkr1Δ:PKR1-2* with H99 at 10.3% (357/3674). We also tested the role of *PKA1* and *PKR1* using the galactose-inducible and glucose-repressible versions of *PKA1* and *PKR1* mutants [6]. In these mutants, when incubated in galactose minimal medium (Fig 9C), the genes are turned on whereas when incubated in glucose minimal medium the genes are turned off (Fig 9D). In galactose minimal medium, PGAL7::PKA1 and PGAL7::PKR1 had titan cell production ratios of 5.6±1.1 and 0.6±0.5 compared to H99, respectively. The proportion of titan cells was significantly increased upon *PKA1* induction [46.9% (794/1724)] and reduced by *PKR1* induction [4.7% (222/5031)] compared with H99 in galactose minimal medium at 14.8% (402/3108) (p<0.0001, Fig 9C). In glucose minimal medium, PGAL7::*PKA1* and PGAL7::*PKR1* had titan cell production ratios of 0.0±0.0 and 2.4±1.0, respectively. The proportion of titan cells was significantly decreased upon *PKA1* repression [0.2% (6/4156)] and increased by *PKR1* repression [25.4%(527/2425)] compared with H99 at 10.3% (357/3674) (p<0.0001, Fig 9D).

**Fig 9.**
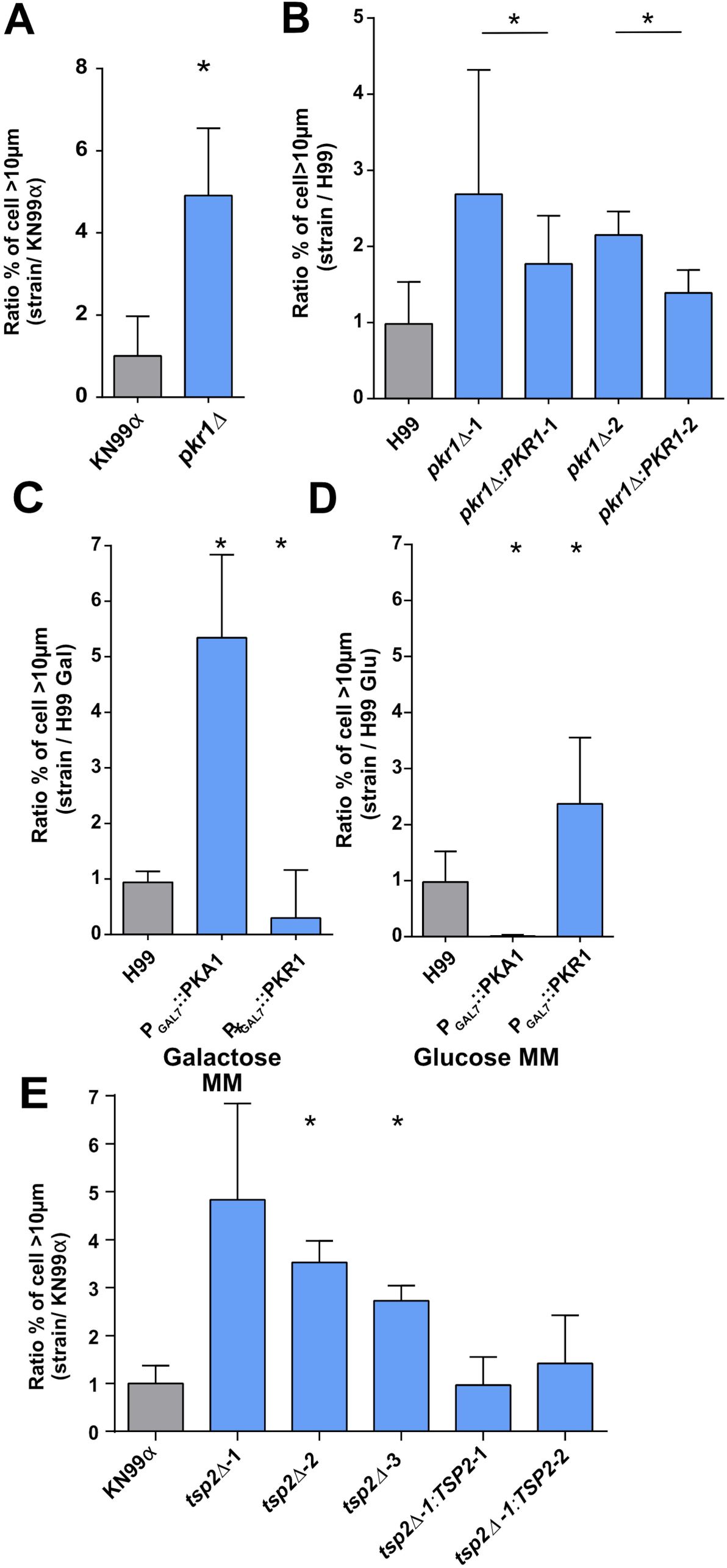
*In vitro* titan cells generation is dependent on the negative regulator *PKR1* and *TSP2*. (**A**) *PKR1* is a repressor of titan cells formation in KN99α background and (**B**) in H99 background. (**C**) in galactose mimimal medium (galactose MM), GAL7 promoter upstream of the *PKA1* and *PKR1* genes induced an increased titan cells formation for PKA1 and a decreased titan cells formation for PKR1 (**D**) In minimal medium with glucose (glucose MM)), GAL7 promoter induced a repression of titan cells formation for PKA1 and an induction of titan cells formation for PKR1. (**E**) Tetraspanin 2 (*TSP2*) is a repressor of titan cells formation. Complementation of the deletion mutants restored the phenotype of KN99α. The ratio to H99 or KN99α, used as a calibrator in each experiment, was calculated for each strain (panel A to E) and results expressed as mean ± SD. Experiments A to G were done in triplicate. To compare the experimental conditions to H99O, Khi2 analysis was performed (*p<0.0001 vs control H99 or KN99α or appropriate mutants)

At least 2 other genes (*TSP2*, *USV101*) have been linked to titan cells formation *in vivo* although their role is less clear, so we directly explored their phenotypes in our *in vitro* protocol. To examine the role of tetraspanin 2 (Tsp2) in titan cell formation *in vitro*, we analyzed three independent tetraspanin 2 deletion mutants (*tsp2Δ-1, tsp2Δ-2* and *tsp2Δ-3*) and two complemented mutants (*tsp2Δ-1:TSP2-1* and *tsp2Δ-1:TSP2-2*) for their titan cell formation compared to the wild-type KN99α. Titan cell production was significantly increased in the deletion mutants (23.9% (518/2161) for *tsp2Δ-1*, 55.7% (990/1778) for *tsp2Δ-2*, 44.3% (714/1610) *tsp2Δ-3*, 4.6% (119/2576)) compared to the wild-type and complemented strains (*tsp2Δ-1:TSP2-1*, 5.8% (228/3877) for *tsp2Δ-1:TSP2-2* and 4.9% (173/3472) for KN99α) (p<0.0001) (Fig 9E). Similarly, *usv101Δ* median cell size (10.6 µm [8.7-12.7]) was higher than the parental strain KN99α (7.6µm [6.6-8.8]) and the proportion of titan cells was 62.7% (1260/2008) for *usv101Δ* vs 10.5% (238/2262) for KN99α (p<0.0001) (S9 Fig).

We finally selected additional sequenced clinical isolates that harbored mutations leading to Usv101 or Cac1 truncation. Bt88 showed a truncation of Usv101 and an increased titan cells formation ratio 0.6±0.4 whereas the other isolates belonging to the VNBII lineage harboring a *CAC1* mutation (Bt133 and Bt31, Bt40, Bt89 and Bt105) did not show increased titan cells formation (ratio 0.0±0.0) (S13 Fig).

## Discussion

We identified and validated a new protocol allowing robust generation of titan cells *in vitro*. This protocol was discovered serendipitously while testing conditions that could induce dormancy in *C. neoformans* [4]. We observed yeast cell enlargement under defined growth conditions, then optimized those conditions for titan cell production. It is important to note that the utility of other published protocols to generate titan cells *in vitro* are hindered by issues with inter-laboratory reproducibility [13, 31]. To establish the inter-laboratory transferability of our protocol, we independently tested it in two other laboratories (K. Nielsen and A. Casadevall) and observed that the protocol produced similar results in all laboratories, although slight variations of the materials and equipment used produced subtle variability in the percentage of titan cells generated in the 3 labs. Interestingly, titan cell production *in vitro* was also optimized by Ballou et al. 2017 and Zaragoza et al. 2017, using a different set of growth conditions [40, 41]. Exploration of the similarities and differences between these protocols will likely identify the critical environmental conditions that trigger titan cell production *in vivo*.

Titan cells obtained *in vitro* exhibited many of the characteristics of *in vivo* titan cells recovered from the lungs of infected mice [17]. Similar to previous work on *in vivo* titan cells, we defined the *in vitro* titan cells as having a cell body size > 10 µm and typical cells with a cell body size ≤ 10 µm [12,13,18,21]. Titan cells generated with our *in vitro* protocol were also polyploids, as previously shown *in vivo* [12, 13]. Melanization was increased in *in vitro* titan compared to typical cells. Capsule size was slightly increased in the *in vitro* titan cells compared to typical cells, but this difference was lower than previously shown *in vivo* [12, 13]. We also demonstrated that, regardless of the capsule size differences between *in vitro* and *in vivo* titan cells, the structure of the *in vitro* titan cell capsule was different to that in typical cells, a phenomenon also observed *in vivo* [13, 22]. The cell wall was thicker in *in vitro* titan cells compared to typical cells, as previously analyzed *in vivo* [22,31,42]. The increased chitin in the titan cell wall results in a detrimental immune response that exacerbates disease[42]. These findings suggest a fundamental difference in titan and typical surface structure that may contribute to reduced titan cell phagocytosis [12,13,21]. These cell surface differences also underscore the complex regulation of these major virulence factors and shows intricate adaptation of *C. neoformans* to both *in vitro* and *in vivo* conditions.

Titan cell generation *in vitro* allowed a detailed kinetic analysis that revealed titan cells are formed between 4 and 8 h. Using calcofluor white staining to follow cell fate and cell division [4, 30], we showed that titan cells were exclusively derived from cells present in the initial inoculum that evolved progressively toward the titan cell phenotype. In contrast, typical cells were a mixture of cells from the initial inoculum and new cell replication. Titan cell division produced typical sized haploid cells, as shown previously for *in vivo* titan cells [12, 13] and confirmed in our study. We published *in vivo* data that validate this observation [4]: using yeasts recovered from the lung of mice at one week after inoculation of *C. neoformans* stained with calcofluor and multispectral imaging flow cytometry, we showed that the Calcofluor^High^ population was associated with yeast cells harboring high cell size parameters compatible with titan cells [4]. *In vitro*, titan cell generation coincided with the appearance of a large vacuole in the yeast cells at 4 and 8 h of incubation. Recent evidence places vacuoles at the center of networks enabling nutrient resources to be degraded, sorted and redistributed [43]. As the vacuole volume occupies much of the total volume of the titan cell body, one can imagine that cell-cycle regulation could be impacted, ultimately leading to polyploidy [42].

The fact that our *in vitro* protocol consistently produced titan cells also allowed us to test factors that influenced their appearance. In terms of environmental factors, we showed that titan cell production was influenced by pre-culture medium, initial pH, light exposure, temperature, type of medium and hypoxia. A metabolic switch between YPD (rich medium) pre-culture and minimal medium (poor medium) incubation was a key factor to induce titan cell generation. This switch is a stress that induces many metabolic modifications and has been studied extensively in *Saccharomyces cerevisiae* [44]. Hypoxia is another stress factor encountered by human pathogenic fungi during infection [45], and a strong signal for titan cell production *in vitro*. Oxygen levels in healthy human tissues are 20–70 mmHg (2.5–9% O_2_), but can be less than 10 mmHg (∼1% O_2_) in hypoxic or inflamed tissues or inside granulomas [46]. We know from previous work on pulmonary aspergillosis that hypoxia has been observed in infected lungs of mice [47]. Titan cells have been reported in human pulmonary cryptococcosis and well-studied in murine pulmonary infection following inhalation [13,17–21], although they are also observed in mouse lungs after intravenous inoculation of animals [4]. These observations lead us to hypothesize that low oxygen levels in the lungs could be a signal for titan cell formation. A major transcriptional regulator of the fungal hypoxia response is the sterol regulatory element-binding protein (Srebp) [48]. Deletion mutants of the *SREBP* gene (*sre1Δ*) in *C. neoformans* display defects in adaptation to hypoxia, ergosterol synthesis, susceptibility to triazole antifungal drugs and cause a reduction of virulence [48]. Importantly, the *sre1Δ* mutant also showed defects in titan cell production *in vitro*, highlighting the role of hypoxia in titan cell production.

Interestingly, quorum sensing is also involved in titan cell production. Indeed, the initial concentration of yeasts in minimal medium dramatically impacted titan cell generation, with 10^6^ cells/mL being the optimal cell concentration to generate titan cells *in vitro*. No titan cell formation was observed with a starting concentration of 10^7^ cells/mL, likely due to rapid consumption of the nutrients preventing metabolic modifications needed to generate titan cells. Alternatively, addition of the quorum Qsp1 peptide [33, 49] to wild type cultures (already producing Qsp1) decreased titan cell production, suggesting that Qsp1 negatively regulates titan cell production. Cleavage and internalization of Qsp1 were critical because the *qsp1Δ, pqp1Δ*, *opt1Δ* mutants all showed increased titan cell production. Pantothenic acid (vitamin B5) has also been implicated in quorum sensing in *C. neoformans* [29]. Addition of pantothenic acid dramatically increased titan cell formation at concentrations between 0.125 and 12.5 µM. These results demonstrate that intercellular communication is important for titan cell formation and that this sensing process involves cell-cell communication instead of simple nutrient sensing.

Induction of titan cells due to the presence of host factors such as temperature, addition of lipids, presence of serum, and antibodies was also tested. The presence of E1 [50] and 18B7 [51] anti-capsular IgG antibodies decreased titan cell production. This decrease could be related to changes in yeast metabolism induced directly by anti-capsular antibodies, as shown previously [52], and provides a new mechanism by which antibodies could alter the course of infection. One could imagine that specific anti-capsular antibodies may not reach cryptococcal cells in the alveolar space at sufficient concentration to impair enlargement. The presence of surfactant protein-D, considered as an opsonin in the lung, could impair antibody fixation [53], thus inhibiting the inhibitory effect of IgG antibodies and allowing titan cell formation in the lung. Interestingly, addition of serum (5% FCS) or L-α-phosphatidylcholine decreased titan cell production, which is different from other protocols for titan cells generation [40, 41]. This difference may be because our protocol induces titan cells through parallel or independent pathways to those triggered by serum or lipids. Overall, these results imply the existence of numerous triggers for titan cell formation mediated through independent signaling pathways. How these pathways ultimately interact, both positively and negatively, to regulate titan cell production still needs to be explored.

Titan cell production was inhibited by addition of fluconazole, flucytosine and cycloheximide – even at concentrations that or below the MIC of the drug. Fluconazole is known to inhibit the 14-alpha-demethylase (Erg11) involved in ergosterol synthesis, leading to plasma membrane instability and accumulation of toxic precursors [54]. Flucytosine is a base analogue leading to inhibition of DNA replication and protein synthesis [55]. Cycloheximide is known to impact protein synthesis through inhibition of translation [27]. Thus, titan cell production likely involves an active process requiring protein and nucleic acid production, as well as plasma membrane integrity (normal ergosterol quantity). Conversely, serial passage in the presence of fluconazole increased titan cell production, suggesting compensatory changes in response to fluconazole also impacted titan cell production. These data have profound implications for *in vivo* titan cell production, as prolonged drug therapy could prevent or enhance titan cell formation. In previous studies, exposure of titan cells to fluconazole selected for aneuploidy and drug resistance in the daughter cells [20]. In contrast, our studies show exposure to cell-wall stress, induced by serial passage on CFW agar, decreased titan cell production. In these sub-culture experiments, we did not investigate subsequent genomic or metabolic changes that arise under these stress conditions. Serial sub-culture could have induced genetic rearrangements (aneuploidy, SNPs, indels) or epigenetic variation that altered titan cell production.

Our protocol is easy to implement for study of the molecular and genetic mechanisms underlying titan cell generation. Our *in vitro* assay allowed us to identify host, environmental and yeast factors that impact titan cell production. By taking advantage of strains harboring genetic differences and clinically relevant genetic truncations, we were able to assess genetic factors modulating titan cell production. However, these studies also highlight that variability in titan cell formation cannot be completely explained by the acquisition of genetic events, with H99-derivative strains showing diversity in titan cell production that does not fully correlate with genetic modifications. The observation that titan cell formation in KN99α differs *in vitro* (lower than H99O) and *in vivo* (equivalent to H99O) highlights this issue and suggests further strain adaptation that are yet to be characterized.

We uncovered new genes involved as positive or negative regulators of titan cell production. Sgf29, is a component of the SAGA complex that binds H3K4me2/3 and recruits histone deacetylases in *S. cerevisae* [56]. The *LMP1* gene is known to be involved in virulence in a mouse model and in mating [34]. We showed here that both genes are positive regulators of titan cell formation, although their mechanism of action remains unclear. We also showed *in vitro* the critical role of the Gpr/PKA/Rim101 pathway in titan cell formation, previously characterized *in vivo* [18]. Gpr5 signals through Gpa1 to trigger the PKA pathway that activates the transcription factor Rim101 [18, 36]. In addition, we identified three genes that are negative regulators of titan cell formation, including *PKR1* (known to act as a regulatory subunit in the PKA pathway [23], *TSP2* that encodes a glucose repressor of laccase in *C. neoformans* [57], and *USV101* that is a pleiotropic transcription factor in *C. neoformans* known to regulate capsule formation and pathogenesis [58].

In *Saccharomyces cerevisiae*, the Pka1/Pkr1 complex is a heterotetramer with 2 catalytic subunits and 2 regulatory units. This complex is dissociated in the presence of cAMP [59]. Moreover, the architecture of the functional domains of *Pkr1* include one interaction domain/dimerization at amino acids 2 to 40 and two cAMP binding domains at amino acids 219 to 351 and 353 to 473, based on INTERPRO data (Fig 8D). Consequently, the cAMP binding domain on Pkr1 is critical for the dissociation of the PKA1/PKR1 complex. Analysis of differences in titan cell production, combined with complete genome sequencing, allowed us to identify naturally occurring mutations in the *PKR1* gene that impact titan cell production. Both *PKR1* mutations have a stop codon (Gly125fs for AD2-06 and Asp14fs for Bt58) that reduces the protein length. Interestingly, AD2-06a was the incident clinical isolate and a recurrent isolate recovered after 13 days of amphotericin treatment (AD2-07) did not harbor this PKR1 mutation. In addition, a *PKR1* mutation leading to intron retention was found in a relapse isolate [60] and shown to be associated with less virulence than the incident isolate. Whether the virulence differences observed with the relapse isolates are linked to titan cell formation needs to be further investigated.

We identified *TSP2* as a negative regulator of titan cell formation based on deletion mutants and complemented strains. *TSP2* is known to interact with the cAMP/PKA pathway – *tsp2Δ* mutant strains phenotype are reversed by the addition of cAMP [57]. These data suggest that *TSP2* inhibits the cAMP pathway and reinforces the major role of cAMP in titan cell formation. No natural *TSP2* mutants were observed in our collection of clinical isolates.

Interestingly, all isolates from the VNBII have a mutation in the *CAC1* gene leading to the functional defect of the Cac1 protein. Out of the six VNBII isolates (S3 Table), only Bt88 that harbored an additional functional abolition of Usv101 was able to produce as much titan cells as H99O did. Therefore, in Bt88, titan cell formation resulted in the equilibrium between the abolition of CAC1 (positive regulator) and USV101 (negative regulator).

Altogether, these results show proof of concept that our *in vitro* protocol can be used to identify and characterize genes required for titan cell production. Our preliminary analysis only identified a handful of genes involved in titan cell production, but it is likely that many more are involved in generation of this complex cell morphology. Our data provide new insights into the genesis of titan cells and the environmental, host and genetic factors that influence their production. Finally, our data show that this *in vitro* protocol can be used to reproducibly generate titan cells that have similar characteristics to titan cells generated *in vivo*. The conditions identified for titan cell formation provide a robust system that could be invaluable to dissect the molecular mechanisms that underlie titan cell formation and allow the identification of naturally occurring mutations that regulate titan cell formation. These studies will enhance our understanding of the impact and mechanisms of yeast morphological changes on pathobiology.

## Material and methods

### Ethics statement

Mice (purchased from Jackson Laboratories, Bar Harbor, ME) were handled in accordance with guidelines defined by the University of Minnesota Animal Care and Use Committee (IACUC), under approved protocol numbers 1010A91133 and 130830852, and in accordance with the protocols approved by JHSPH IACUC protocol M015H134. All animal experiments were performed in concordance with the Animal Welfare Act, United States federal law, and NIH guidelines.

### Strains and culture medium

The strains and clinical isolates of *C. neoformans* used in the study are listed in S4 Table. The study was started with H99 strain called H99O that was kindly provided by J. Heitman (Duke University, Durham, NC) in the late 90’s. The reference strain KN99α and strains from the Madhani collection were provided from Kirsten Nielsen’s lab and the Fungal Genetic Stock Center [61], respectively. *C. neoformans* strains were grown in liquid Yeast Peptone Dextrose (YPD, 1% yeast extract (BD Difco, Le Pont de Claix, France) 2% peptone (BD Difco), 2% D-glucose (Sigma, Saint Louis, Minnesota, USA)) and in minimal medium (MM, 15mM D-glucose (Sigma), 10 mM MgSO4 (Sigma), 29.4mM KH2PO4 (Sigma), 13mM Glycine (Sigma), 3.0 µM Thiamine (Sigma), [32]). Minimum inhibitory concentration (MIC) of H99O for fluconazole (FLC) and flucytosine (5FC) (both purchased from lsachim, Shimadzu Group Company, Illkirch-Graffenstaden, France) were determined by the EUCAST method and were 8 and 4 mg/L, respectively.

### *In vitro* protocol for titan cells generation

*C. neoformans* strain from stock cultures stored in 20% glycerol at −80°C was cultured on Sabouraud agar plate at room temperature (step 1). After 2 to 5 d of culture, approximately 10^7^ cells were suspended in 10 mL YPD in a T25cm^3^ flask and cultured 22 h at 30°C, 150 rpm with lateral shaking until stationary phase (final concentration=2×10^8^cells/mL) (step 2). Then, one mL of the suspension was washed twice with MM. The cell concentration was adjusted to 10^6^ cells/mL in MM and the suspension was incubated in a 1.5 mL tube (Eppendorf) with the cap closed, at 30°C, 800 rpm for 5 d using an Eppendorf Thermomixer (Hamburg, Germany) (step 3). Cell size was determined as described below. Cells with body size >10 µm were considered as titan cells as described [12]. Results are expressed as median cell size [interquartile range, IQR] or as median [IQR] of the proportion of titan cells in a given condition for H99 or as a ratio compared to the proportion of titan cells obtained with the H99O in experiments involving other strains (clinical isolates, other H99 strains and mutants). In specific experiments, 10^4^ cells/mL were incubated in 100 well plate (Fischer Scientific) and incubated at 30°C with agitation in the Bioscreen apparatus (Fischer Scientific).

In specific experiment using P_GAL7_ inducible mutants in H99, MM with galactose at 15mM (galactose MM) was used in parallel to MM containing glucose (see above).

### Capsule size analysis

Yeasts were observed after India ink staining and capsule thickness was determined as the size of the thickness in pixel of the white area surrounding the cell wall imaged with an Olympus AX 70 microscope and analyzed using the ImageJ software available at https://imagej.nih.gov/ij/ and the Multi_measures plugin.

### Chitin content and capsule structure quantification

Multispectral flow cytometry was used to quantify chitin content after calcofluor white staining (CFW, fluorescent brightener 28, 0.0001 µg/mL CFW in PBS) and capsule structure after immunostaining of three anti-capsular antibodies (E1 IgG1 monoclonal antibody [50], both 2D10 [29] and 13F1 [29] IgM monoclonal antibody 30 m at 10 µg/mL) and then incubation with FITC coupled anti-IgG or -IgM secondary antibodies (15 m at 1:1000 concentration in PBS). The antibody 18B7 has not been used for this specific experiment because it produces aggregation that prevented ImagestreamX testing. Pictures were taken in flow and analyzed using various existing algorithms. We used ImageStreamX with the INSPIRE software (Amnis Corporation). Cell suspensions were adjusted to 10^7^ in 200 µL and 10,000 cells were recorded at 40-fold magnification in 3 different channels including the bright field channel (BF) and 2 fluorescence channels (channel 1: 430-505nm [CALCO]; channel 2: 470-560nm [Anticapsular antibodies]). Data analysis was performed using the IDEAS software (Amnis Corporation) after fluorescence compensation procedures. The first step consists in the definition of a mask that delineates the relevant pixels in each picture. Then, 54 algorithms (calculations made for each event within a defined mask) are available to analyze size, texture, location, shape or signal strength. Using basic algorithms, unfocused events, yeasts aggregates were excluded [4]. First titan cells (TC) and typical cells (tC) were selected based on a dot plot Area/Diameter. We decided to avoid overlap between populations and select well separated population based on their size after control using the bar added on the picture of the yeasts (see Fig 2A-2B). For chitin content, the calcofluor intensity histogram using the Intensity of algorithm in channel 01 have been generated for TC and tC (see Fig 2C-2D). For capsule structure, the algorithms dedicated to structure analysis were tested and Modulation and Bright details intensity R7 algorithms in channel 2 have been found to separate the capsule structure of titan cells from typical cells populations. For each population of interest, the geometric mean was calculated using the IDEAS software.

Additional experiments using fluorescence microscopy for chitin content measurement was performed after calcofluor white staining (CFW, fluorescent brightener 28) adapted from [42]. Briefly, 10^7^ *C. neoformans* cells in 10 mL MM were washed once and 500 µL of 3.7% formaldehyde in PBS was added. Cells were incubated at room temperature for 30-40 m, inverting the tube every 5 m. Samples were washed twice in PBS, cell concentration was adjusted to 10^6^ cell/mL. The supernatant was removed and 1 mL of 0.0001 µg/mL CFW in PBS was added, and incubated 5 m at 25°C. Cells were then washed twice in PBS. Results were expressed as median [IQR] of the mean fluorescence intensity/pixel/cell after picture analysis as described below.

### Capsule immunofluorescence and melanization analysis

Capsule immunofluorescence (IF) of titan cells was done by incubating approximately 5×10^6^ cells/mL with 10µg/mL of murine-derived monoclonal antibodies to the cryptococcal capsule (IgG1 18B7, IgG1 E1, IgM 12A1, IgM 2D10 [29, 50] in blocking solution (1% bovine serum albumin in PBS). Cells and mAb mixtures were done in 1.5 mL microcentrifuge tubes at 37 °C for 1 h under continuous mixing. Next, cells were washed three times with PBS by centrifugation (5,000 rpm for 5 m at room temperature) and incubated for 1 h at 37 °C with 5 µg/mL fluorescently labelled secondary-mAbs, goat anti-mouse IgG1-FITCs or IgM-TRITCs (Southern Biotech) in blocking solution and 1 µg/mL of Uvitex2b (Polysciences, Warrington, PA) solution to visualize the fungal cell wall. Cells were washed three times with PBS by centrifugation, mounted in glass coverslips and imaged with an Olympus AX 70 microscope equipped with blue, green and red fluorescent filters using 40x and/or oil immersion 100x objectives. Capsule immunofluorescence of titan cells preparations performed in two independent experiments gave consistent results.

Titan cells melanization was induced following step 3. Cells were washed once with minimal medium, suspended in 1mL of minimal medium supplemented with 1mM of L-DOPA (Sigma D9628), transferred to a 5mL Erlenmeyer flask (for normal oxygenation) and incubated at 30 °C under continuous mixing at 200 rpms for 3 d. Since melanin is resistant to acid hydrolysis, a spherical melanin “ghost” remains following incubation of black cells with 12N HCl for 1 h at 100 °C (a reduced and modified version of the procedure in [62]. Acid-resistant melanin “ghosts” were washed three times in PBS by centrifugation and visualized using light microscopy.

Melanization was measured using imageJ in Icy software by manually circling each cell and measuring the mean gray intensity / pixel /cell. Blackness was calculated as the maximum mean grey intensity minus the mean grey intensity / pixel /cell. Increasing melanin content will result in higher blackness.

### N-acetylglucosamine quantification

H99 cells were grown in MM for 48 h *in vitro*. Cells were centrifuged 2 m at 14,000 rpm and were then washed twice with sterile water. These cells were exposed to γ-radiation to remove layers of the capsule polysaccharide [13]. Cells resuspended in sterile water were transferred to a 24-well flat-bottom plate and irradiated for 45 m: dose 560 Gy (56,000 rad). titan cells and tyical cells were separated [20]. Washed irradiated cells were filtered using CellMicroSieves (BioDesign Inc. of New York, Carmel, NY) with a 10 µm pore size. The CellMicroSieves were rinsed with PBS to remove typical cells from the filter. To recover the titan cells population, the CellMicroSieves were inverted and the membrane was washed with PBS. The TCs population was concentrated by centrifugation at 12,000 g for 1 m. To recover the typical cells population, the filter flow-through was concentrated by centrifugation at 12,000 g for 1 m.

Cellular chitin quantification was adapted from [63]. Purified *in vitro* titan cells and typical cells were collected by centrifugation at 14,000 rpm for 2 m and the media were removed. Dry weights were measured following 2-3 d of evaporation at 37°C. Dried pellets were extracted with 1 mL 6% KOH at 80°C for 90 m. Samples were centrifuged at 14,000 rpm for 20 m. Each pellet was suspended in 1 mL PBS and spun again. Each pellet was suspended in 0.2 mL of McIlvaine’s Buffer (0.2 M Na2HPO4, 0.1 M citric acid, pH 6.0). Five µL of purified *Streptomyces griseus* chitinase (5 mg/mL in PBS) was added to hydrolyze chitin to Glu-cNAc and incubated for 3 d at 37°C. Chitinase-treated samples were spun at 14,000 rpm for 1 m, each 10 μL of sample supernatant was combined with 10 μL 0.27 M sodium borate, pH 9.0. Samples were heated to 99.9°C for 10 m. Upon cooling to room temperature, 100 μl of DMAB solution (Ehrlich’s reagent, 10 g p-dimethylaminobenzaldehyde in 12.5 mL concentrated HCl, and 87.5 mL glacial acetic acid) was added, followed by 20 m incubation at 37°C. Hundred µL was transferred to 96-well plates, and absorbance at 585 nm was recorded. Standard curves were prepared from stocks of 0.2 to 2.0 mM of Gluc-NAc (Sigma, Saint Louis, Missouri, USA). The amount of Gluc-NAc was calculated as mmol/g cells (dry weight). Results are expressed as median [IQR].

### DNA content measurement

A 96-well microtiter plate was filled with 200 µL of a 10^6^/mL cell suspension in PBS and centrifuged 5 m at 4000 rpm. The pellet was suspended in 150 µL of ethanol 70% and incubated in the dark 1 h at 4°C. After discarding the supernatant, a 50 µL mix composed of 44 µL NS (0.01M Tris HCL pH 7.2, 1 mM EDTA, 1mM CaCl2, 0.25 M Sucrose, 2.12 mM MgCl2, 0.1 mM ZnCl2), 5µL RNase A at 10mg/mL and 1.25µL PI at 0.5mg/mL was added in each well as described [64]. After a 30 m incubation at 30°C in the dark, the plate was sonicated 1 m and each sample diluted at 1:40 in 50mM Tris HCl. The fluorescence intensity was measured using the Guava easyCyte 12HT Benchtop Flow Cytometer (Guava, MERCK, Kenilworth, New Jersey). Selection of singlets by gating allowed (i) determination of PI intensity on channel YelB (583/26) in FSC/SSC^high^ (TC) and FSC/SSC^low^ (tC); (ii) determination of the FSC/SSC distribution in PI^high^ and PI^low^ population. FlowJo software v.10 was used to analyze the data. The graphs of the number of yeasts were normalized to the mode to depict the data in terms of ‘% of max’. The % of max denotes the number of cells in each bin (the numerical ranges for the parameter on the *x* axis) divided by the number of cells in the bin that contains the largest number of cells.

### Determination of the ancestry of titan cells by flow cytometry

Knowing that CFW staining does not alter *C. neoformans* viability and that daughter cells harbored lower CFW signal due to partial cell wall transmission from mother to daughter cells [4, 30], we analyzed cell size and CFW fluorescence intensity of the progenies of titan cells and typical cells following the *in vitro* protocol on 10^6^ cells of H99O pre-stained with CFW (channel BluV 448/50 using Guava).

### Dynamic imaging

Budding rates were determined after yeasts (10^5^ cells composed of titan cells and typical cells) previously incubated using our protocol or *in vivo* (see below) were directly deposited in a 35 mm sterile culture dish in minimal medium without agitation and incubated at 30°C. Pictures were taken every 2 or 5 m by phase microscopy using the Axiovert 200M inverted microscope with 40X or 20X objectives (Carl Zeiss MicroImaging, NY), used in conjunction with an AxiocamMR camera.

Cell size evolution over time was assessed for strain AD2-06a by dynamic imaging (Nikon Biostation). Briefly, 35 mm sterile culture dish (Hi-Q4, Nikon) were coated for 1 m with E1 antibody at 2 mg/L in order to provide anchor for the capsule. Yeasts (10^5^ cells) were added in 1 mL MM and incubated at 32°C for 18 h. Series of 221 images were taken by phase-contrast microscopy every 5 m at ×100 magnification. Merging was done using ImageJ software in Icy Software.

### Impact of various factors on titan cells generation

To analyze the various factors that could impact titan cells generation, we modified the various steps of our *in vitro* protocol. For step1, stress was produced by 8 subcultures (twice a week for one month) on agar medium or on agar supplemented with CFW (20mg/L) or with fluconazole (32mg/L). For step 2, the pH of MM (normally at 5.5) was set at 4, 7 or 8.5 without buffering. For step 3, initial cell concentration (from 10^4^ to 10^7^cells/mL) was tested. Hypoxia was generated physically by closing the cap of the Eppendorf tube during 5 d or chemically upon incubation in MM supplemented with 1 nM CoCl_2_, cap closed, as already described [65]. The production of titan cells was also assessed in the presence of various reagents added at step 3: (1) Qsp1 peptide (NFGAPGGAYPW, [33]) (Biomatik, Cambridge, Canada) was resuspended at 10mM in water and stored at −80°C until use at 10 µM final with the scrambled peptide (AYAPWFGNPG) as a control; (2) pantothenic acid purchased from Sigma (Saint-Louis, Missouri, USA) used at 125 µM; (3) monoclonal anti-capsular antibodies E1 [50] and 18B7 [51] used at a final concentration of 166 µg/mL in MM [66]; (4) decomplemented fetal calf serum (FCS, Invitrogen, Carlsbad, CA, USA) at 5 % in MM; (5) L-α-Phosphatidylcholine from egg yolk (Sigma, Saint-Louis, Missouri, USA) was extemporaneously reconstituted at 5 mM in MM; (6) antifungal drugs (fluconazole and flucytosine) were tested at the concentrations close to the MIC (2-fold dilutions) with the diluent (DMSO or water) as control. Results are expressed as median [IQR]. Growth in the presence or in the absence of antifungal drugs was evaluated by enumeration of yeast cells concentration at step 4 of our protocol using the Guava cytometer, starting from 10^6^ cells inoculated at step 1.

### Production and isolation of titan cells from infected mice

*C. neoformans* strains were cultured overnight at 30°C in YPD broth medium (BD, Hercules, Canada). Yeast cells were collected by centrifugation, washed with phosphate buffered saline PBS and resuspended in sterile saline. For titan cells analysis *in vivo*, groups of 6- to 8-week-old C57BL/6J mice (Jackson Labs, Bar Harbor, Maine) were anesthetized by 5% isofurane inhalation for 1-5 m, infected intranasally with 2 × 10^5^ cells in a 40 µL volume and sacrificed at D6. In these experiments, 84% of titan cells were obtained. For mutant screening, groups of 6- to 8-week-old C57BL/6J mice (Jackson Labs, Bar Harbor, Maine) were anesthetized by intraperitoneal pentobarbital injection and infected intranasally with 5 × 10^6^ cells in a 50 µL volume. Infected mice were sacrificed by CO_2_ inhalation at 3 d post-infection. The lungs were harvested, homogenized, and then resuspended in 10 mL PBS supplemented with collagenase (1 mg/mL) [13]. Cell homogenates were incubated for 1 hour at 37°C with agitation, and washed several times with double distilled water. The *C. neoformans* cells were fixed with 3.7 % formaldehyde for 40 m, washed 3 times with sterile PBS, and then resuspended in sterile PBS. The proportion of titan cells and typical cells were determined by microscopy. Data presented were from 3 mice per strain, except for strains *sgf29Δ* in H99O, H99S that had 2 mice per strain.

### Mutant generation

We PCR amplified the *PKR1* and *TPS2* genes using the primer KN99α DNA as substrate and the following primers (PKR1F: AAGCTTggaatgaagatgaaattagtacgtg; PKR1R: ACTAGTgtccatcattgctgtaacttggttg; TSP2F: GAGCTCaactccgatgatcatggactcgg; TSP2R: GAGCTCtgcccaagagactagagtgtaacc). The 2559 bp TPS2 and the 2000 bp PKR1 amplicons were cloned in the pGEMT easy vector (Clontech) and sequenced. The pNE609 and pNE610 plasmids were then constructed by cloning the *PKR1* and *TPS2* DNA fragments into the pSDMA57 plasmid [67] using the SpeI/HindIII and SacI cloning sites, respectively.

To create transformants, the plasmids pSDMA57 containing PKR1 amplicon was linearized with BaeI and biolistically transformed into AD2-06a and Bt156 clinical strains. To complement the *tsp2Δ* mutant, pSDMA57 plasmid containing *TSP2* gene was linearized with BaeI and biolistically transformed into the *tsp2Δ* mutant strain. All transformants were selected on YPD supplemented with neomycin. Genomic DNA was purified from the transformants and PCR was used to check the presence of *PKR1* and *TSP2* genes in the transformed strains. PCR reactions contained 1 μl gDNA, 2.5 μl of each of the 10 mM primer stocks (PKR1 forward, PKR1 reverse, TSP2 forward, TSP2 reverse) 5 μl Taq buffer, 4 μl dNTPs, 0.25 μl ExTaq polymerase (New England Biolabs, USA) and 34.75 μl sterile water. The cycling parameters were 35 cycles of 94°C for 20 seconds, 54°C for 20 seconds and 72°C for 90 seconds. Products were visualized using electrophoresis with 0.8% TAE agarose gel. To differentiate between random integration, single insertion, and tandem insertion into the safe haven, we performed a similar PCR as above using primers UQ1768, UQ2962, UQ2963, and UQ3348 as previously described [67].

### DNA Sequencing, variant identification, and bioinformatic analysis

Genomic DNA was adapted for Illumina sequencing using Nextera reagents. Libraries were sequenced on an Illumina HiSeq to generate 101 base reads. most data was previously described [39, 68] and one additional isolate was newly sequenced for this study (AD2-07) (NCBI SRA accession SRR5989089). Reads were aligned to the *C. neoformans* H99 assembly (GenBank accession GCA_000149245.2 [34] using BWA-MEM version 0.7.12 [69]. Variants were then identified using GATK version 3.4 [70], where indels were locally realigned, haplotypeCaller was invoked in GVCF mode with ploidy = 1, and genotypeGVCFs was used to predict variants in each strain. All VCFs were then combined and sites were filtered using variantFiltration with QD < 2.0, FS > 60.0, and MQ < 40.0. Individual genotypes were then filtered if the minimum genotype quality < 50, percent alternate allele < 0.8, or depth < 5. Variants were then functionally annotated with SnpEff version 4.2 [71]. For phylogenetic analysis, the 535,968 sites with an unambiguous SNP in at least one isolate and with ambiguity in at most 10% of isolates were concatenated; insertions or deletions at these sites were treated as ambiguous to maintain the alignment. Phylogenetic trees were estimated using RAxML version 8.2.4 [72] under the GTRCAT model in rapid bootstrapping mode. For determination of Pkr1 architecture domains, the INTERPRO tool was used (http://www.ebi.ac.uk/interpro/protein/J9VH50).

### Pictures analysis using Icy software and statistical analysis

To increase the number of events analyzed in each condition tested/each parameter analyzed (cell size, capsule size and chitin content), pictures of 3-5 fields were taken with an AxioCam MRm camera (Carl Zeiss, Oberkochen) at x40 on interferential contrast microscope (DMLB2 microscope; Leica, Oberkochen). Image were then analyzed (for cell size and chitin content) using Icy software v.1.9.2.1.[73] (icy.bioimageanalysis.org) and a specific plugin (HK-Means plugin (http://icy.bioimageanalysis.org/plugin/HK-Means) that allows analysis of multiple structures from a bright field. Preliminary experiments were done to compare results obtained with Icy to “manual” measurements by analyzing about 200 cells on the same pictures for 3 independent experiments. In subsequent experiments, results were pooled for a given condition from 2 to 3 independent experiments after good reproducibility was assessed. Statistical analysis was performed with STATA software (College Station, Texas, v13.0). To validate the cell size determination using the Icy software, the intraclass correlation coefficient was calculated. The ability of the automated method to classify the *C. neoformans* cells as titan cells or typical cells compared to visual measurement was evaluated using the Kappa test [74]. To compare titan cells generation in the various conditions, non-parametric tests were performed using the Kruskal-Wallis test for multiple comparisons or Mann Whitney test as required. GraphPad Prism software (v.6) was used to generate graphs.

### Accession numbers

All sequence data from this study have been submitted to NCBI BioProject (https://www.ncbi.nlm.nih.gov/bioproject) under accession number PRJNA174567.

The AD2-07 sequence is available in the NCBI SRA under the accession number SRR5989089 (https://www.ncbi.nlm.nih.gov/sra/SRR5989089/)

## Acknowledgments

We warmly thank Pr James Kronstad and Melissa Caza for sending us their collection of deletion and complemented mutants of the *PKR1* gene, as well as their PGal:PKR1 and PGal:PKA conditional mutants. We thank Dr Tihana Bicanic, Shichina Kannmbath, Charles Giamberardino and Jennifer Tenor who kindly provided specific clinical isolates. The authors want to thank Marie Desnos-Ollivier for her help with Sanger sequencing, JL Tinevez for technical assistance in Biostation experiments, Stéphane Dallongeville for help with Icy Software, Stevenn Volant for biostatistics, Pierre-Henri Commere for FACS analysis, Frederique Moyrand for performing PCR and plasmid preparations. We thank Quigly Dragotakes for helping us determining cell and capsule sizes in specific experiments. The authors acknowledge Hiten Madhani and members of his laboratory for the gene deletion collection that has been made available ahead of publication to the scientific community.

## Supporting informations

**S1 Fig. In vitro protocol of titan cells generation.** The protocol followed four steps: (1) *C*. *neoformans* H99O from a frozen stock culture at −80°C was cultured on Sabouraud agar for 2-5 d; (2) Approximately 10^7^ yeasts were then suspended in 10mL of liquid Yeast Peptone dextrose (YPD) and incubated under agitation (150 rpm) at 30°C for 22 h (stationary phase); (3) 1 mL of the culture was then washed twice in minimal medium (MM), then 10^6^ yeasts were resuspended in 1mL of minimal medium (MM) Ph5.5, in a 1.5 mL Eppendorf tube and incubated at 800 rpm for up to 120 h using an Eppendorf thermomixer; (4) A mixture of typical cells and of titan cells was ready for analysis.

**S2 Fig. Among yeasts recovered at the end of the *in vitro* protocol, those with the highest DNA content have the biggest cell size.** DNA content was analyzed after propidium iodide (PI) staining of yeast cells obtained at the end of our protocol (H99O induced), in a control haploid strain (H99O cultured in Sabouraud agar, H99O-sab) and in a control diploid strain (AD7-77 cultured in Sabouraud agar). Part of the population of H99O-induced had a higher PI (blue arrow) fluorescence intensity than the haploid control (upper panel). Gating on the PI intensity showed that the increase in the PI fluorescence intensity from <20K to >40K corresponded to increase in cell size (FSC) (red arrows) compared to the diploid (AD7-77) and haploid (H99O Sab) control (lower panel).

**S3 Fig. The FSC^high^/CFW^high^ population of yeasts correspond to titan cells (TC).** Cells obtained using our *in vitro* protocol were stained with CFW and sorted by flux cytometry according to size (FSC) and CFW fluorescence intensity (left panel). Sorted yeasts were observed using bright field and fluorescence microscopy (right panel) (bar=10µm). Typical cells (tC) were FSC^low^/CFW^low^.

**S4 Fig. Chitin characterization and melanization of titan cells.** (**A**) Chitin was denser in titan cells (TC) than in typical cells (tC) according to CFW fluorescence intensity/pixel/cell measured by Icy software after CFW staining (0.01 µg/mL) at step 4 of the protocol (*p<0.0001). Dots represent individual cells, and boxes median and IQR for 400 cells each (*p<0.001, pooled measurements from 3 independent experiments). (**B**) N-acetylglucosamine (GlcNAc), the monomer component of chitin, was increased in titan cells (TC) compared to typical cells (tC) *in vitro* (left panel) and *in vivo* (right panel) as measured by a biochemical method after gamma-irradiation of the yeasts to remove the capsule, allowing a better separation of titan cells and typical cells. Each dot represents result from independent experiments (n=7). Results are presented as median and IQR (p<0.001). (**C**) Comparing the blackness of the cell body of titan cells (TC) and typical cells (tC) upon melanization conditions showed that titan cells contained more melanin than typical cells. (Bar=10µm). (**D**) Melanization was more important in titan cells (TC) than typical cells (tC) (*p<0.0001) based on the calculation of the max – mean grey value/pixel of each melanin ghost measured (n=19 for titan cells and n=531 for typical cells) using the ImageJ in Icy software. Each dot represents an individual cells and boxes median and IQR.

**S5 Fig. Capsule structure of titan cells.** (**A**) Using multispectral flow cytometry and capsule staining using anticapsular monoclonal antibodies (mAb), we discriminated the distribution of titan cells and typical cells with almost no overlap between both population with 2D10 mAb *in vitro* and *in vivo*, based on the algorithm modulation and Bright details intensity R7. Overlap in the staining characteristics of titan cells and typical cells were observed for E1 (IgG1) and 13F1 (IgM) antibodies. (**B**) Immunofluorescence staining with the anti-capsular mAbs 2D10, 12A1, 18B7, and E1 does not uncover major differences in capsular structures between titan cells (white arrows) and typical cells (black arrows). Each panel correspond to the same cells observed after staining with (a) India ink; (b) calcofluor white; (c) one of the Mabs; (d) merge from c and d. (bar=10µm).

**S6 Fig. Growth is maintained in the presence of antifungals after our protocol.** 10^6^ cell/mL were inoculated at step 1 of our protocol, then cell growth was evaluated by enumerating cell concentration obtained at step 4 of our protocol using Guava flow cytometer for cell counting. (**A**) Compared to control, fluconazole did not modify cell growth in MM whereas (**B**) flucytosine (5FC) reduced it, when used at concentration near the minimum inhibitory concentration (MIC) for 5 d (*p<0.0001 compared to unexposed control). The fluconazole and flucytosine MICs for H99O were 8 mg/L and 4 mg/L, respectively. Experiments were done in triplicates (bars represent mean ± SD). (**C**) Cycloheximide also reduced cell growth at 0.0001 and inhibit cell growth at 0.001 mg/mL (*p<0.0001 compared to unexposed control)

**S7 Fig. Test of iterative subcultures affect titan cells formation with or without the presence of active molecules (CFW or fluconazole)** Step 1 was modified by sub-culturing H99O 8 times over one month on Sabouraud agar alone (Sub8), or supplemented with 20mg/L CFW (Sub8+CFW) or with 32mg/L fluconazole (Sub8+FLC). Compared to initial culture (0Sub), 8 sub-cultures (8Sub) decreased significantly the cell size (** p<0.0001, vs 0Sub control). In addition, iterative subcultures on CFW and FLC decreased and increased significantly the cell size compared to the 8Sub control, respectively. Median and IQR are shown in black for each condition (* p<0.0001 vs 8Sub control). The numbers above each condition represent the proportion of titan cells observed. The experiments were performed in triplicate and pooled (mean cell counted ± SD = 2455±913).

**S8 Fig. Titan cells generation is dependent on various genes *in vivo*.** (**A**) Strains from the H99 lineage harbored variable abilities to generate titan cells with H99O and KN99α *in vivo* compared to the other H99 strains (S, L, W, CMO18). (**B**) The *sgf29*Δ and *lmp1*Δ mutant strains show a decrease in titan cells generation in various H99 backgrounds *in vivo* compared to H99O and rescued by complementation. Each experiment was done in triplicates. Results are presented as stacked bar of the proportion of titan cells (titan cells) and regulars cells (typical cells), * p<0.0001 vs control H99O.

**S9 Fig. The proportion of titan cells generated is dependent on various genes and requires signaling through the Gpr/PKA/Rim101 pathway.** The different H99 strains harbored variable abilities to produce titan cells compared to H99O *in vitro* (**A**) and in vivo (**B**). *Sgf29Δ* and *lmp1Δ* deletion mutants show a decrease in titan cells generation in various H99 backgrounds compared to H99O *in vitro* (**C**) and in vivo (**D**). Complementation in strains *lmp1Δ:LMP1* and *sgf29Δ:SGF29* in H99S background restored the phenotype of H99S. (**E**) Rim101 and *GPR4* and *GPR5* and *CAC1* are required for titan cells generation *in vitro* in H99 and KN99α. The ratio to the value obtained for H99O used as a calibrator in each experiment was calculated for each strain. Bar represent mean ± SD (mean cell counted=600). Khi2 test was performed to compare the experimental conditions to H99O, they were performed in triplicates and pooled (*p<0.0001, **p<0.0001, when the comparison was done with the parental strain H99S.)

**S10 Fig. *USV101* and *SRE1* deletion influenced titan cells formation** (**A**) *usv101Δ* is a repressor of titan cells formation. (**B**) The *sre1Δ* mutant strain decreased titan cells formation compared to the parental strain KN99α. The ratio to KN99α, used as a calibrator in each experiment, was calculated for each strain and results expressed as mean ±SD. To compare the experimental conditions to KN99α, Khi2 analysis was performed (*p<0.0001).

**S11 Fig. Chr9 ploidy does not influence titan cells generation.** A panel of 7 clinical isolates with partial Chromosome 9 duplication (left panel) was tested for its ability to generate titan cells. Only H99O and AD2-06a exhibited increased cell sizes (middle panel). The proportion of titan cells was 67.9 % (431/667) for AD2-06a, 32.1% (429/1339) for H99O, and 4.2% (51/1227), for Ug2459 (Khi2 compared to H99O, *p<0.0001) with the ratio of the proportion of titan cells to that produced in H99O shown in the right panel.

**S12 Fig. *PKR1* mutations influence median cell size** Strains with *Pkr1* loss of function mutation showed a variable ability to produce titan cells depending on the resulting truncated proteins. The clinical isolate AD2-07 which did not harbor the *PKR1* mutation was recovered from the CSF of an HIV-positive patient on d 13 of amphotericin B treatment while AD2-06a was recovered from its initial CSF. The median cell size (5.9 µm [5.2-6.6]) was significantly decreased in AD2-07 and increased in AD2-06a (8.5 µm [7.0-13.0]) compared to H99O (7.7 µm [6.6-9.2]) (p<0.0001). Except Bt156 (median of 7.7 µm [6.4-9.4]), the others strains had a significantly decreased median size compared to H99O (p<0.0001), 6.9 µm [6.0-7.8] for 8-1 strain; 6.5 µm [5.8-7.1] for Bt77, 6.4 µm [5.7-7.2] for Bt117; and 5.4 µm [4.7-6.1] for Ug2462. Experiments were done in triplicate and pooled.

**S13 Fig. Non-synonymous mutation in *USV101* enhances titan cells generation based on clinical isolates analysis.** Bt88 harbored a truncated Usv101 protein due to a frameshift mutation. The titan cells generation is negative in Bt31, Bt40, Bt89, Bt105 and Bt133 and increased for Bt88 with a ratio at 0.6±0.4 and a proportion of titan cells of 21.6% (423/1958) compared to H99O 38.5% (729/1890). Experiments were done in triplicate and pooled.

**S1 Movie. Time lapse imaging of titan cells and typical cells generated *in vitro* (after 5 d) allowed to grow.** Titan cells produced normal sized daughter cells upon incubation in fresh MM (at 30°C, one picture every 2min during 24 h).

**S2 Movie. Time lapse imaging of titan cells and typical cells generated *in vivo* (6 d post infection, intranasal route) allowed to grow.** Titan cells produced normal sized daughter cells upon incubation in fresh MM (at 30°C, one picture every 5 m during 24 h at ×400 magnification using transmitted light (white bar=10 µm, NC = typical cells).

**S3 Movie. Time lapse imaging showing mothers cells increasing in time allowing titan cells generation first produced between 8 and 12 h of incubation.** Dynamic imaging of yeasts from the AD2-06a *C. neoformans* clinical isolate using the Nikon Biostation IM. Yeasts were seeded in MM on a culture dish that was previously coated with E1 at 2 mg/L. Images were taken every 5 m for 24 h at ×100 magnification using transmitted light (white bar=10 µm)

**S1 Table.** Gene disrupted in AD2-06a but not in closely related isolated AD3-55a or AD3-41a

**S2 Table.** Clinical isolates with chromosome 9 ploidy variation

**S3 Table.** Strains harboring Pkr1 loss-of-function mutations used in this study

**S4 Table.** Strains used in this study

